# RNA-induced PRC2 inhibition depends on the sequence of bound RNA

**DOI:** 10.1101/2024.08.29.610323

**Authors:** Jiarui Song, Liqi Yao, Anne R. Gooding, Valentin Thron, Wayne O. Hemphill, Karen Goodrich, Vignesh Kasinath, Thomas R. Cech

## Abstract

Methyltransferase PRC2 (Polycomb Repressive Complex 2) deposits histone H3K27 trimethylation to establish and maintain epigenetic gene silencing. PRC2 is precisely regulated by accessory proteins, histone post-translational modifications and, particularly, RNA. Research on PRC2-associated RNA has mostly focused on the tight-binding G-quadruplex (G4) RNAs, which inhibit PRC2 enzymatic activity in vitro and in cells, a mechanism explained by our recent cryo-EM structure showing G4 RNA-mediated PRC2 dimerization. However, PRC2 binds a wide variety of RNA sequences, and it remains unclear how diverse RNAs beyond G4 associate with and regulate PRC2. Here, we show that variations in RNA sequence elicit distinct effects on PRC2 function. A single-stranded G-rich RNA and an atypical G4 structure called a pUG-fold mediate PRC2 dimerization nearly identical to that induced by G4 RNA. In contrast, pyrimidine-rich RNAs, including a motif identified by CLIPseq in cells, do not induce PRC2 dimerization and instead bind PRC2 monomers with retention of methyltransferase activity. Only RNAs that dimerize PRC2 compete with nucleosome binding and inhibit PRC2 methyltransferase activity.

CRISPR-dCas9 was adapted to localize different RNA elements onto a PRC2-targeted gene, revealing RNA sequence specificity for PRC2 regulation in cells. Thus, PRC2 binds many different RNAs with similar affinity, however, the functional effect on enzymatic activity depends entirely on the sequence of the bound RNA, a conclusion potentially applicable to any RNA-binding protein with a large transcriptome.

## INTRODUCTION

In the last decade, non-canonical RNA-binding proteins that do not contain conventional RNA-recognition or RNA-binding motifs have been of increasing interest. Examples include histone modifiers ^1,2^, chromatin architecture remodelers ^3–5^, DNA methylases ^6,7^, transcription factors ^8–10^ and metabolic proteins ^11,12^, many of which participate in tuning differential expression of genes. The binding of RNA has been reported to have various functions, including promoting complex recruitment and enhancing target recognition specificity as a positive regulator and, on the other hand, inhibiting enzymatic activity and limiting target accessibility as a negative regulator, emphasizing the under-characterized but critical role of RNA in epigenetic regulation.

The histone methyltransferase Polycomb Repressive Complex 2 (PRC2) is a prevailing model system for studying the mechanism and significance of RNA binding to epigenetic modifiers ^13–15^. PRC2 trimethylates lysine 27 of histone H3 (H3K27me3), which is a repressive mark for gene expression and essential for normal development and cell differentiation ^16,17^. Studies have shown that RNAs capable of folding into G-quadruplex structures (G4 RNAs) bind PRC2 to inhibit its activity in cells ^18–21^, and our recent cryo-EM structure of a G4 RNA-bound PRC2 complex revealed the molecular mechanism of such inhibition ^22^. Instead of blocking PRC2-chromatin binding by simple steric inhibition, the G4 RNA bridges two PRC2 protomers to form a dimer that specifically buries PRC2 amino acids required for docking the histone H3 tail and binding the nucleosomal DNA.

However, the PRC2 interactome has been reported to contain thousands of pre-mRNAs and lncRNAs, and it includes pyrimidine-rich RNAs and G-rich RNAs with fewer than four G-tracts, which cannot fold into a G4 structure ^20,23–30^. These other RNAs differ from canonical G4 structures in nucleotide composition, shape, charge distribution, and availability of H-bond donors/acceptors, raising the questions of how PRC2 recognizes various RNAs and, more importantly, what the functional consequences are for PRC2 associating with these different RNAs.

Here, we use biochemical methods and electron microscopy (EM) to investigate PRC2 binding to diverse RNAs. Notably, RNA-mediated PRC2 dimerization is sequence-dependent. G-rich RNAs, including single-stranded RNA, can induce PRC2 dimerization. Flexible loops of PRC2 accommodate different RNA structures, showing how G-rich RNA beyond G4 can inhibit PRC2. In contrast, PRC2 binds pyrimidine-rich RNAs as a monomer, which does not prevent PRC2 activity in vitro or in cells. Overall, our study expands RNA-mediated PRC2 regulation to multiple RNA elements, enhancing the understanding of how diverse RNAs can mediate different functional consequences for this key regulator of epigenetic gene silencing.

## RESULTS

### PRC2 complexes bind multiple RNA sequences

Although PRC2-RNA binding has been studied previously ^29,31–34^, we sought to extend these analyses by examining a series of PRC2 complexes with a more complete set of RNAs involving different sequences and structural features. We first prepared the six-subunit PRC2 complex with accessory proteins AEBP2 and JARID2, because accessory proteins have previously been implicated in RNA binding ^18,35^. To this end, seven 50-nucleotide RNA oligos were synthesized (Fig. 1A). TERRA contains four repeats of the human telomeric sequence and can form a single G4 structure, mimicking the noncoding RNA transcribed from telomeres ^36^ and shown to recruit PRC2 to telomeres ^37^. TERRA_mut_ is a single-stranded G-rich RNA that has the same base composition as TERRA, but the sequence is mutated to eliminate consecutive guanines, preventing G4 formation. The pUG-fold RNA has 20 repeats of GU, which forms an atypical RNA quadruplex and distinct from the canonical G4 by having three G quartets and one additional U quartet ^38^ (Figure S1B). Poly(A) does not bind to the five-subunit PRC2 ^29^ and serves as a negative control in our characterization of the six-subunit complex. Finally, Poly(U), Poly(C), and P10 are pyrimidine-rich RNAs. In particular, P10 has a C and U repetitive sequence, which was initially identified as a PRC2 binding consensus in cells by crosslinking and immunoprecipitation (CLIP) ^30^. Native gel electrophoresis confirmed the formation of RNA secondary structures in TERRA and pUG-fold RNAs (Fig. S1A).

**Figure 1.**
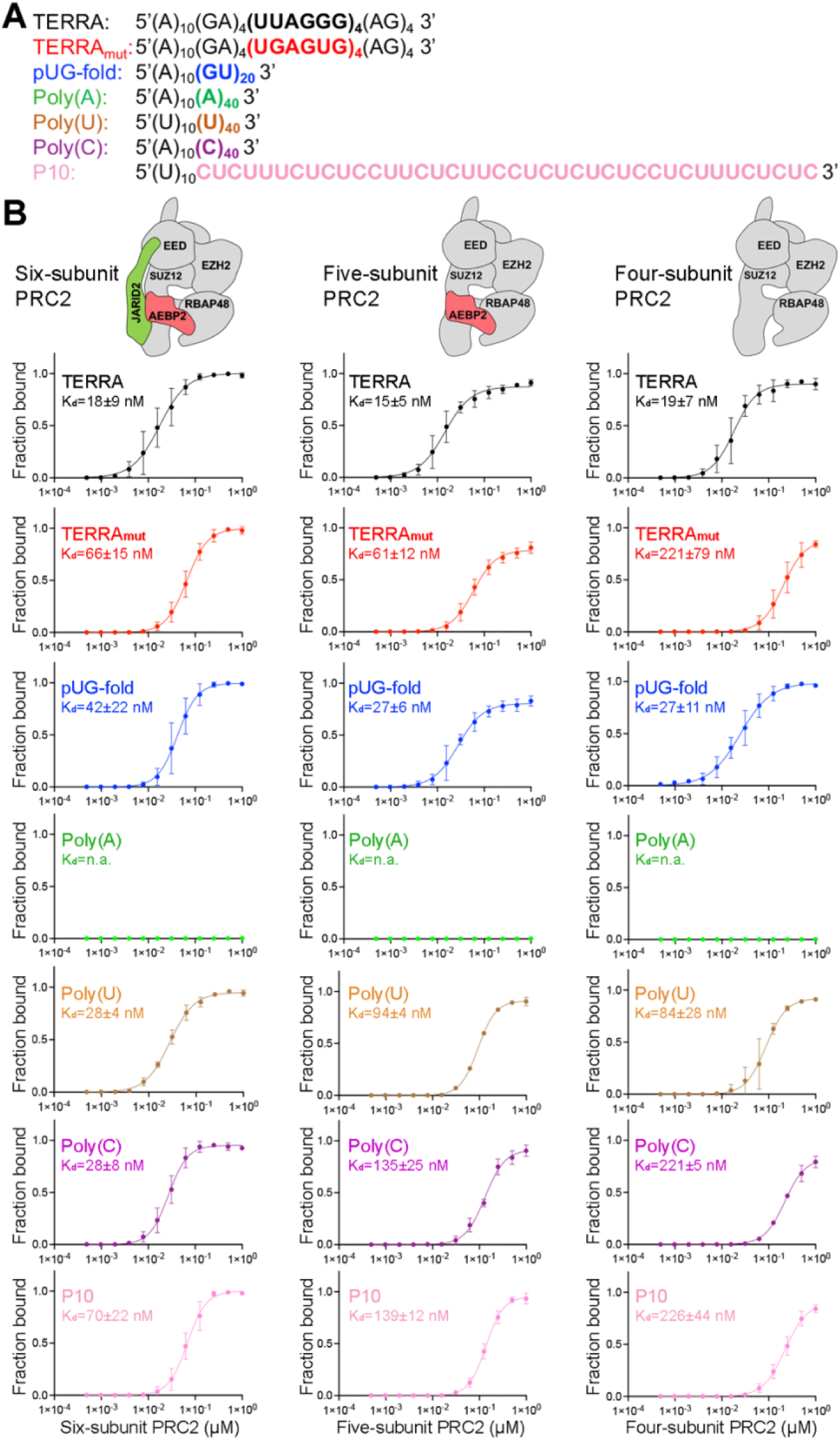
PRC2 complexes bind RNAs of various sequences and structures. (A) The seven RNA oligonucleotides used in this study. Colors highlight the functional sequences, while the 5’ (A)_10_ or (U)_10_ linkers provide flexibility for binding to streptavidin EM grids. (B) Top: schematic representation of six-, five- and four-subunit PRC2 used in this study. Bottom: quantification of three EMSA replicates and the corresponding K_d_ values of PRC2 binding RNAs. Error bars are mean ± standard deviation.

Electrophoretic mobility shift assays (EMSA) were performed with PRC2 and various RNAs in 100 mM KCl to approximate nuclear ionic conditions, stabilizing G4 structures (Fig. 1B and Fig. S1D). PRC2 bound TERRA, Poly(U), and Poly(C) with the highest affinity, approximately 2-fold, 3-fold, and 4-fold stronger than pUG-fold, TERRA_mut,_ and P10, respectively. When binding was instead performed in 100 mM LiCl to preclude G4 formation, the three G-rich RNAs bound PRC2 with similar affinities (Fig. S2), indicating that the six-subunit PRC2 complex retained the preference for binding the folded G4 RNA but could also bind unfolded G-rich RNAs. Surprisingly, six-subunit PRC2 also bound single-stranded Poly(U) and Poly(C) with affinities similar to that of folded TERRA (K_d_(TERRA)=18±9 nM, K_d_(PolyU)=28±4 nM, and K_d_(PolyC)=28±8 nM).

Our initial hypothesis for PRC2 preferring G4 was that the G4 RNA has a more condensed electrostatic charge distribution in its compact structure, but the high binding affinities of single-stranded pyrimidine-rich RNAs seen here do not support this idea. We therefore tested whether accessory proteins included in the six-subunit complex were aiding the binding to these additional RNA structures and sequences. The five-subunit PRC2 (missing JARID2) had near identical binding affinities for the three G-rich RNAs as the six-subunit, but it had substantially reduced binding affinities for all pyrimidine-rich RNAs. Interestingly, the four-subunit holoenzyme (missing AEBP2 and JARID2) showed weak binding to TERRA_mut_ but not TERRA or pUG-fold (Fig. 1B and Fig. S1C-F). The K_d_(TERRA_mut_)/K_d_(TERRA) ratio was 12 for the four-subunit PRC2, significantly larger than the ratio of 4 obtained for the six-subunit and five-subunit complexes. Additional reduction of binding affinities to the Poly(C) and P10 RNAs was also found for the four-subunit complex compared to the five-subunit complex.

Thus, our systematic characterizations indicate that the four-subunit holoenzyme contains all essential protein elements to recognize G4 as the most preferred RNA, consistent with our cryo-EM structure showing that the catalytic subunit EZH2 is predominantly responsible for G4 RNA recognition. Alternatively, the six-subunit complex binds RNA sequences and structures more broadly. Accessory proteins significantly elevate PRC2 association of pyrimidine-rich RNAs (by JARID2) and most of single-stranded RNAs (TERRA_mut_, Poly(C), and P10 by AEBP2). More generally, these biochemical results help explain the in vivo results showing a broad PRC2 transcriptome containing many lncRNAs and pre-mRNAs ^20,24,26,27,39,40^.

### PRC2 dimerization is RNA sequence-dependent

We recently adapted the streptavidin-affinity EM grid method ^41^ to allow specific selection and enrichment of ribonucleoprotein (RNP) complexes ^22^. Here, we used 5’-biotinylated RNAs and negative-staining EM to determine the architectures of PRC2 bound to different RNA sequences (Fig. 2A). Reference-free 2D-class averages of PRC2-TERRA, PRC2-TERRA_mut,_ and PRC2-pUG-fold RNPs exhibited PRC2 dimerization with similar protomer arrangement as shown in the cryo-EM structure of the (GGGAA)_4_ G4 RNA-mediated PRC2 dimer ^22^. The finding that TERRA_mut_ could mediate PRC2 dimerization was interesting, given that it bears no resemblance to a folded G4 RNA with respect to shape, hydrogen bond donor/acceptor potential, or electrostatic charge distribution. In contrast, the majority of PRC2 RNP particles remained monomeric when bound to pyrimidine-rich RNAs. We validated that the affinity grid method captured only RNA-bound complexes, as Poly(A) RNA did not capture any recognizable particle (Fig. S3). Quantifying two negative-staining EM replicates of each PRC2-RNA complex revealed that dimerized PRC2 consistently comprised 70% to 90% of all recognizable particles when incubated with G-rich RNAs and less than 20% with pyrimidine-rich RNAs (Fig. 2B).

**Figure 2.**
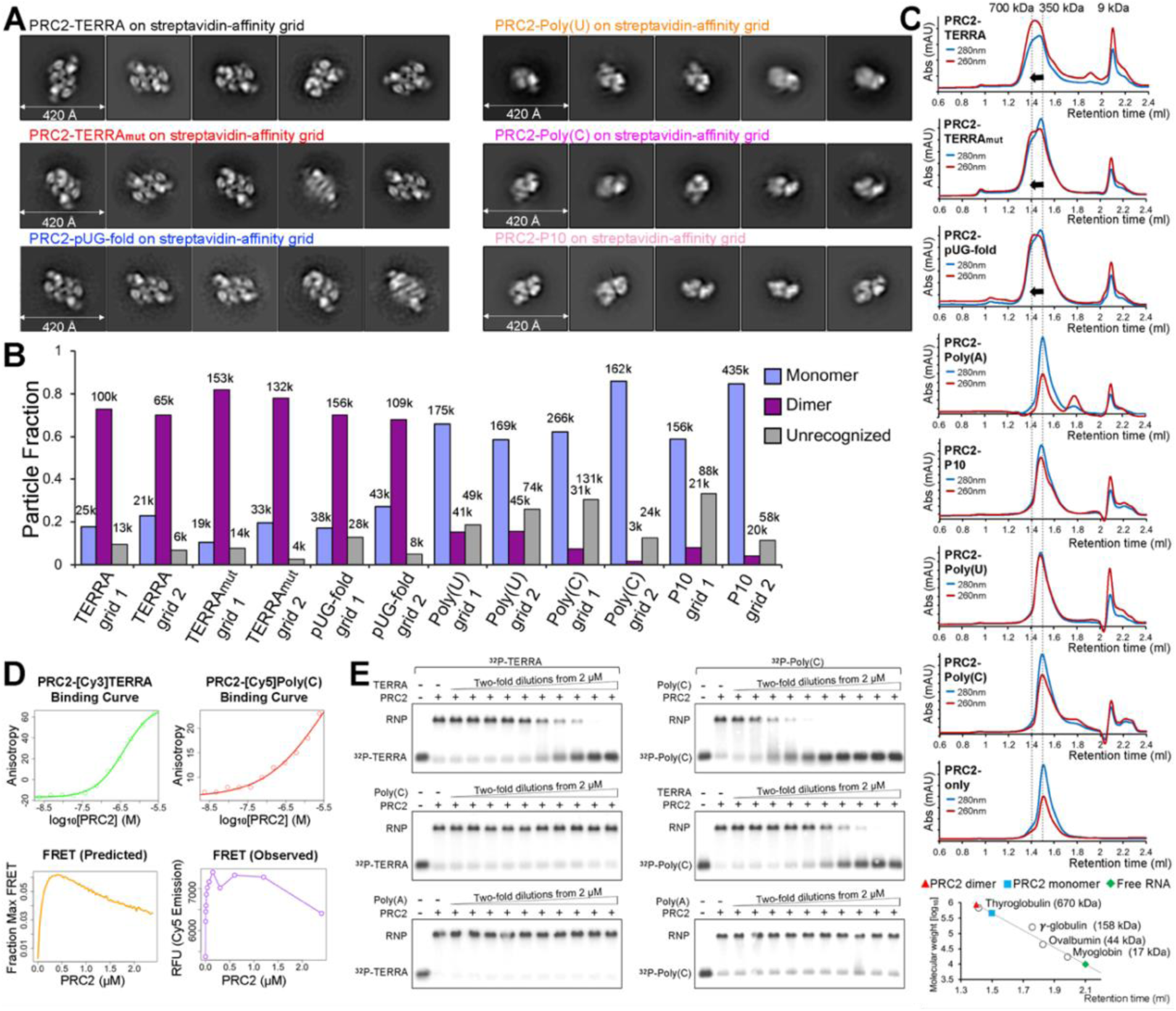
RNA sequence-dependent dimerization of PRC2 and evidence that G4 and Poly(C) RNAs bind to different sites on PRC2. (A) Negative-staining EM provided 2D-class averages of six PRC2-RNA complexes, three RNAs inducing PRC2 dimers and three binding to PRC2 monomers. The 2D-class averages are shown in order of prevalence from left to right. (B) Quantification of two negative-staining EM replicates (two independent EM grids, each grid utilizing PRC2 from a separate preparation). (C) Top: size-exclusion chromatography of PRC2 preincubated with RNAs and mock (protein only). Abs, absorbance; mAU, milli-absorbance unit. Bottom: standard curve used to estimate the molecular weights of complexes. (D) FRET assay indicates the formation of TERRA-Poly(C)-PRC2 ternary complex. Top left: PRC2-TERRA binding measured by fluorescence polarization (FP) of Cy3 signal. Top right: PRC2-Poly(C) binding measured by FP of Cy5 signal. Binding curves for each ligand were fit with Equation 1 (see Methods). Bottom left: Predicted FRET behavior as a function of protein concentration is calculated from the binding curve fits via Equation 2, which assumes the two ligands bind independently. Bottom right: the observed FRET. All data were collected from the same reaction containing fixed, equal concentrations of both [Cy3]-TERRA and [Cy5]-Poly(C) RNAs and increasing concentrations of PRC2. (E) Representative competitive EMSA assays. PRC2 was preincubated with ^32^P-labeled TERRA (left) or Poly(C) (right) before adding competitor RNAs at different concentrations. Successful competition removed ^32^P-labeled RNA from PRC2. Three replicates gave the same conclusion.

Analytical size-exclusion chromatography was performed to validate our EM observations in solution (Fig. 2C). PRC2 alone and PRC2-Poly(A) chromatographed at a molecular weight of approximately 350 kDa, equal to the sum of the individual PRC2 subunits. PRC2-Poly(U), PRC2-Poly(C), and PRC2-P10 eluted at the same retention volume with the 280 nm/260 nm absorbance ratio significantly lower than 2, consistent with PRC2 binding pyrimidine-rich RNAs as a monomer. This chromatography trace was distinct from the profiles of PRC2 and G-rich RNAs, which formed RNPs with a molecular weight of 700 kDa. Our results suggest that RNA nucleotide composition predicts the RNA-mediated dimerization of six-subunit PRC2.

The two distinct RNA sequence-dependent binding modes of PRC2 inspired us to hypothesize that PRC2 could utilize separate protein elements to engage different RNAs. If so, then the PRC2 binding of G-rich and pyrimidine-rich RNAs might not be mutually exclusive. We tested this idea by fluorescence resonance energy transfer (FRET) (Fig. 2D) and RNA-competition EMSA (Fig. 2E). The clear concentration-dependent FRET signal at low PRC2 concentrations indicated the formation of a PRC2-TERRA-Poly(C) tertiary complex. The FRET signal gradually diminished when PRC2 protein increased and the two RNAs began to bind separate PRC2 complexes, suggesting that the two RNA ligands bind independently rather than cooperatively. In RNA-competition EMSA, Poly(C) could not remove the pre-bound radiolabeled TERRA from PRC2 (Fig. 2E left). In the other direction, it required considerably more TERRA than Poly(C) itself to effectively compete with pre-bound Poly(C) (Fig. 2E, right). Both results support the conclusion that competition between different RNA ligands is always less efficient than self-competition. In summary, PRC2 can simultaneously engage G-rich and pyrimidine-rich RNAs, and the sequence of the RNA determines its binding mode with PRC2.

### Cryo-EM structure of TERRA_mut_-bound PRC2 dimer

To understand the interaction of PRC2 with an RNA that does not fold into a canonical G4 structure, we solved a 3.3 Å cryo-EM structure of the single-stranded TERRA_mut_-bound PRC2 (Fig. 3A, Fig. S4 and S5, Table S1). In this structure, the PRC2 dimer had the identical architecture as the G4 RNA-mediated complex, including the protein-protein interface assembled by the CXC domain of the catalytic subunit EZH2. This dimer interface is key to explaining RNA inhibition of PRC2 activity, because it buries amino acids that are required for histone H3 tail and nucleosome binding in the active PRC2 monomer ^42^.

**Figure 3.**
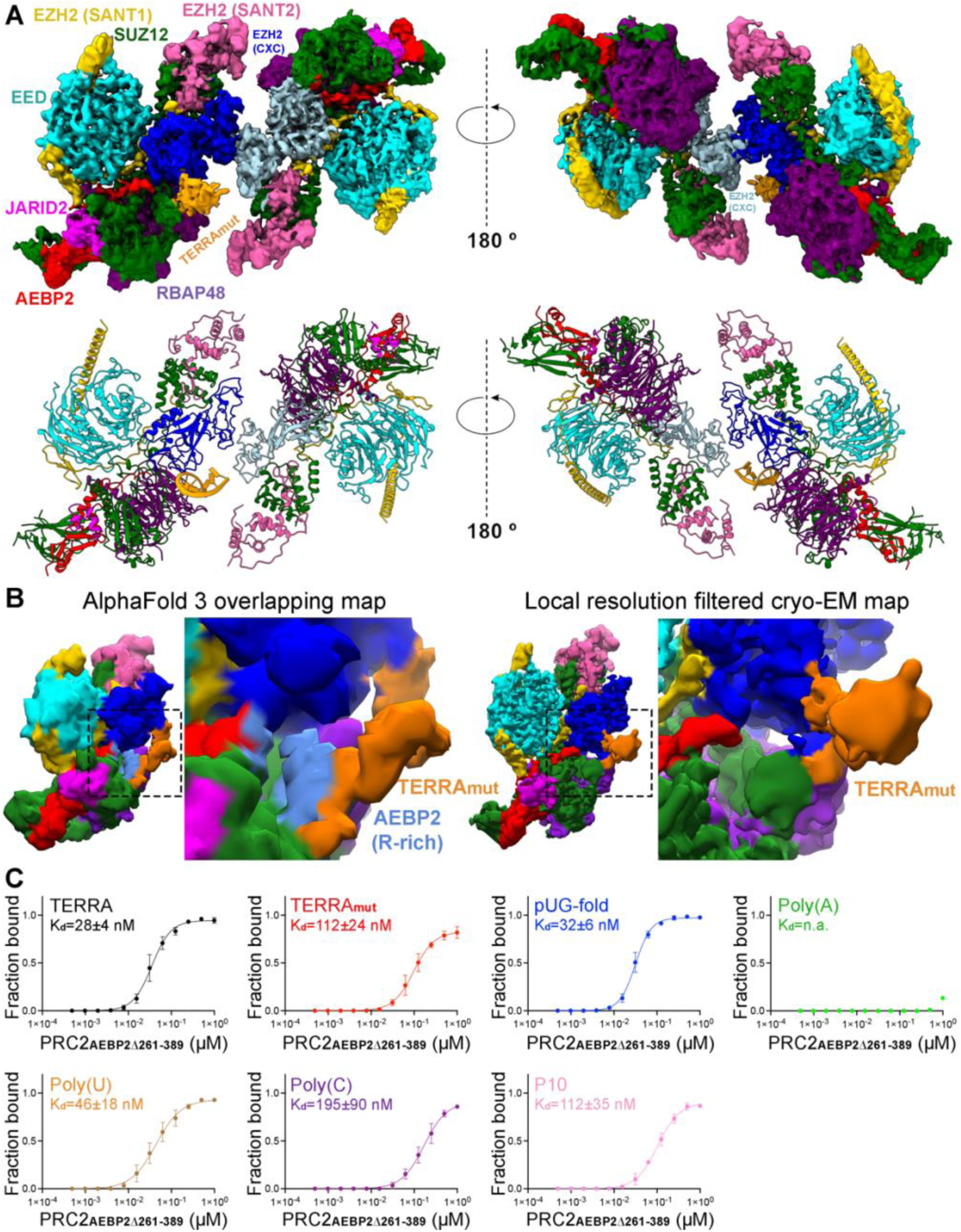
PRC2-TERRA_mut_ structure and accessory protein AEBP2 implicated in contributing to RNA binding. (A) Top: Cryo-EM density map of TERRA_mut_ RNA-associated PRC2 dimer. EZH2 (CXC-SET) of protomer 1 in blue, EZH2 (CXC-SET) of protomer 2 in light blue, and TERRA_mut_ RNA in orange. Bottom: The corresponding atomic model. (B) Left: Density map generated by keeping only the overlapped regions of the top four AlphaFold3 models (also see Fig. S7) (see Methods). Right: Local resolution filtered cryo-EM density from multibody refinement of TERRA_mut_ RNA-associated PRC2. Dashed boxes are zoomed in. The remaining density of TERRA_mut_ after averaging AlphaFold models has similar size and position compared to the observed cryo-EM density. (C) Quantification of three EMSA replicates of six-subunit PRC2_AEBP2Δ261-389_ complex binding different RNAs. Error bars are mean ± standard deviation.

We could confidently identify one TERRA_mut_ RNA density, suggesting a single TERRA_mut_ is sufficient to induce PRC2 dimerization (Fig. 3A). Because PRC2 utilizes flexible loops to engage RNA, allowing multiple RNA orientations and conformations, we could not obtain high-resolution details to model TERRA_mut_ de novo. The location of TERRA_mut_ partially overlapped with the location of G4 RNA from our previous cryo-EM reconstruction (Fig. S6). This can be explained if both PRC2 protomers contribute to RNA binding in a pincer-like arrangement. The G4 and TERRA_mut_ RNAs are suspended at similar positions in the middle of the two PRC2 protomers without strict RNA structure requirement. Also, the TERRA_mut_ density is attached to the bottom lobe of one PRC2 protomer in a region of RBAP48 and AEBP2, which was less structured in the G4 RNA-bound map (Fig. S6). This difference may indicate that the single-stranded TERRA_mut_, expected to be more extended than a G4, interacts with additional protein surfaces. Overall, the use of flexible loops to bind RNA, the formation of a dimer with RNA suspended between the protomers, and the additional protein surfaces available for extended RNA elements appear to provide PRC2 with the potential to accommodate disparate RNA structures.

We also attempted to solve the cryo-EM structure of the pyrimidine-rich RNA-bound PRC2 and failed. A technical limitation of the affinity grid method is the requirement of applying three support layers on the surface of the EM grid: streptavidin crystal, biotinylated lipid, and thin carbon (≍2 nm in thickness) ^41,43^. Thus, most small-sized particles (e.g., monomeric PRC2) do not have a strong contrast of intensity after computational streptavidin-lattice subtraction, insufficient for reliable particle picking and further alignment. Also, monomeric PRC2 might be more susceptible to conformational heterogeneity, as most available monomeric PRC2 structures utilized crosslinking before cryo-EM preparation. For these reasons, we could not obtain a high-resolution structure of the monomeric PRC2 RNP.

### AEBP2 is implicated in contributing to single-stranded RNA binding

Because the details of RNA-protein interaction were not resolved by cryo-EM, we used AlphaFold3 ^44^ to generate models of TERRA_mut_, Poly(U), and Poly(C) bound to a single PRC2 (Fig. S7-S10). In parallel, we performed multibody refinement of our cryo-EM consensus map to focus on the single PRC2 protomer that interacted with TERRA_mut_, which could be directly compared to the AlphaFold3 models. Across all PRC2-TERRA_mut_ predictions, the overall organization of PRC2 subunits remained consistent and agreed well with published PRC2 structures ^45–51^, while TERRA_mut_ RNA bound to the expected region of PRC2 but had variable conformations and orientations with respect to PRC2 in each predicted model (Fig S7, A-D).

AlphaFold3 predictions had high confidence for PRC2 subunits (chain iPTMs=0.4-0.6) and low confidence for TERRA_mut_ (chain iPTM=0.1). Predictions of PRC2-TERRA_mut_, PRC2-Poly(U), and PRC2-Poly(C) showed no significant difference in terms of the overall PRC2 and RNA arrangement and details of protein-RNA interactions.

Despite the low confidence in RNA modeling by AlphaFold, it was gratifying to see that the predicted TERRA_mut_ always overlapped with the density we designated as the RNA in our cryo-EM multibody reconstruction. To better present this feature, we generated a surface map including only regions overlapping between all top AlphaFold models (Fig. 3B and Fig. S7E); this eliminating flexible regions of the RNA, similar to the principle used in cryo-EM reconstruction that averages several hundred thousand particles. This surface map had identical architecture to the actual cryo-EM map, particularly with only part of TERRA_mut_ being visualized, explaining how the intrinsic flexibility of PRC2-RNA binding affects our EM map.

AlphaFold3 indicated the importance of PRC2 accessory protein AEBP2, utilizing zinc-finger motifs and an arginine-rich segment (379-KRRKLKNKRRR-389), for binding all three RNAs (Fig. S8-S10). The second and third C2H2-type zinc-finger motifs within AEBP2 are in the periphery of the RNA, engaging RNA via electrostatic interactions with the phosphate backbone and hydrogen bonds with both backbone and bases. The residues responsible for this interaction differ between models, indicating that, unlike typical zinc-finger motifs that recognize specific DNA sequences, AEBP2 zinc-finger motifs have less strict RNA sequence specificity. In addition, the arginine-rich segment of AEBP2 contains nine positively charged amino acids, showing a strong electrostatic potential to bind many RNAs. To test these predictions, we prepared the six-subunit PRC2 complex with truncated AEBP2 (PRC2_AEBP2Δ261-389_), which lacks both zinc-finger motifs and the arginine-rich segment.

Compared to WT PRC2, PRC2_AEBP2Δ261-389_ had a modest reduction of TERRA_mut_ binding (K_d_(PRC2_WT_)=66±15 nM and K_d_(PRC2_AEBP2Δ261-389_)=112±24 nM, P=0.05) and more dramatic decrease of Poly(C) binding (K_d_(PRC2_WT_)=28±8 nM and K_d_(PRC2_AEBP2Δ261-389_)=195±90 nM, P=0.03) (Fig. 3C and Fig. S11). The binding of all other RNAs was not statistically different between WT and the AEBP2 mutant. This result is largely consistent with the reduced RNA binding affinities we observed for the PRC2 four-subunit holoenzyme (Fig. 1B), in which AEBP2 is absent, indicating that the zinc-finger motifs and arginine-rich segment are primarily responsible for enabling AEBP2 of PRC2 to accommodate diverse RNAs.

### RNAs that induce dimerization inhibit PRC2 activity

Most biochemical characterizations of RNA-mediated PRC2 inhibition have focused on short G4-forming sequences ^18,21,22,34,52^ or long RNAs that likely fold into complex 3D architectures containing multiple structural elements (e.g., HOTAIR and Xist lncRNAs) ^18,20,32,53^. The effect of simple G-tract sequences other than canonical G4 and especially the pyrimidine-rich RNAs on PRC2 activity has not been measured.

Our structures of RNA-induced PRC2 dimers have demonstrated that the EZH2 dimer interface prevents nucleosomal DNA and H3 tail accessibility to the PRC2 catalytic groove. As expected, all G-rich RNAs that can induce PRC2 dimerization inhibited radiolabeled tri-nucleosomes from binding PRC2 in nucleosome-RNA competition assays (Fig. 4A). Pyrimidine-rich RNAs, instead, did not interfere with nucleosome binding, indicating that PRC2 has separate nucleic acid-binding segments specialized for nucleosomal DNA, pyrimidine-rich RNA, and G-rich RNA. Consistent with this model, methyltransferase activity assays of PRC2 with co-incubation of RNAs showed a progressive reduction of H3K27 methylation when we included TERRA, TERRA_mut,_ and pUG-fold in trans, but exhibited no change with pyrimidine-rich RNAs and Poly(A) negative control (Fig. 4B). IC_50_ of TERRA, TERRA_mut_, and pUG-fold were all around 500 nM, and complete inhibition was achieved at higher RNA concentrations. The observed IC_50_ is higher than the K_d_ because the inhibition of PRC2 is determined not only by the K_d_ of the inhibitor but by the RNA-PRC2 stoichiometry ^54^, given that 600 nM PRC2 was used in the assay. Our biochemical characterizations clearly illustrate that different RNA sequences mediate distinct functional consequences in regulating PRC2. Only RNAs that induce dimerization prevent PRC2 binding to nucleosomes and subsequently inhibit PRC2 activity.

**Figure 4.**
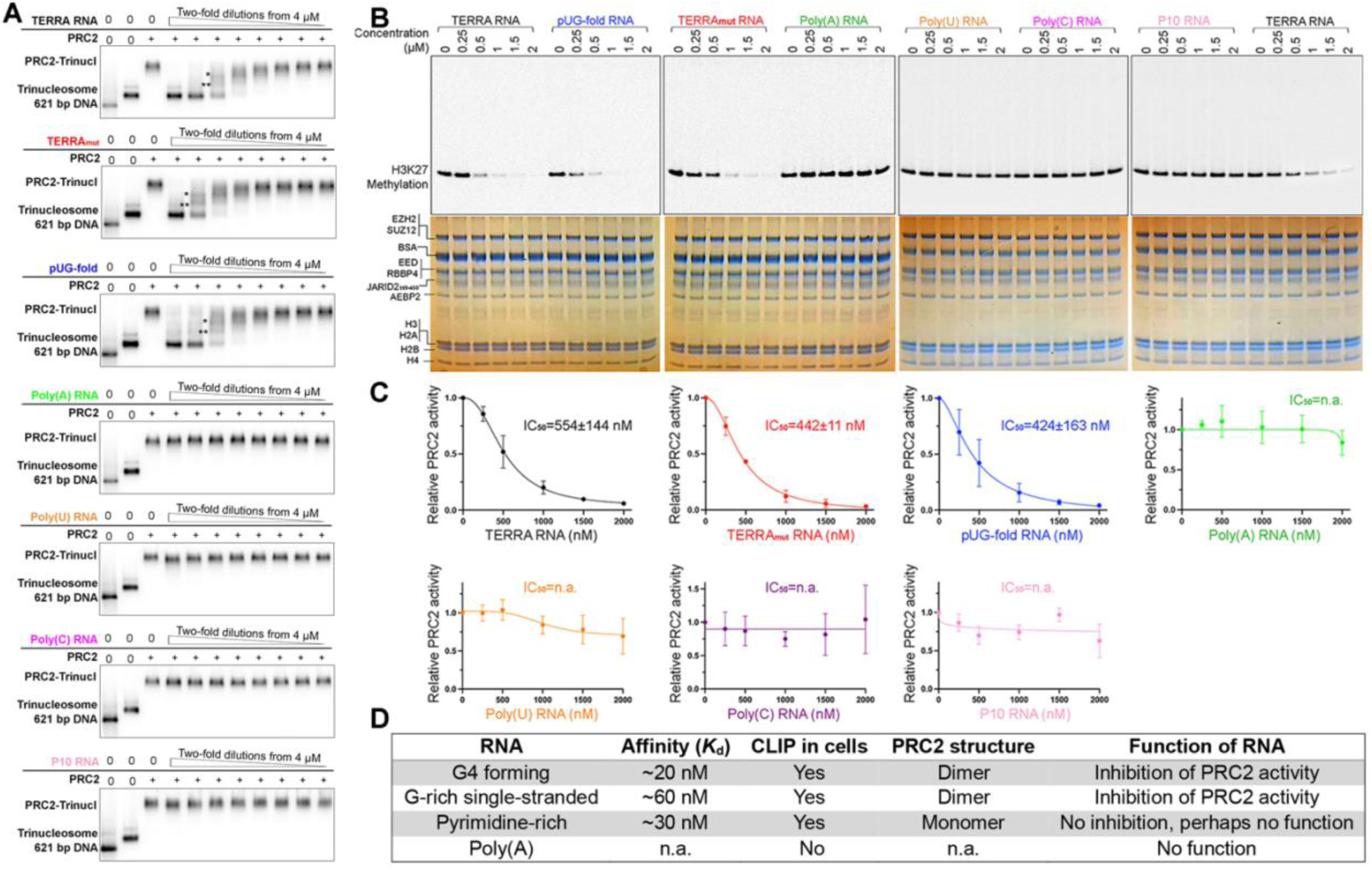
Only RNAs that induce dimerization inhibit PRC2 binding to nucleosomes and activity. (A) Representative EMSA gels of nucleosome-RNA competition assays. PRC2 was preincubated with ^32^P-labeled trinucleosomes before adding competitor RNAs at different concentrations. Successful competitions removed ^32^P-labeled trinucleosomes from PRC2, represented by reduction of PRC2-nucleosome signal and appearance of free trinucleosomes. This experiment was performed three times with equivalent results. Incomplete PRC2-trinucleosome complexes are indicated by * and **. We assume two of three nucleosomes were occupied by PRC2 in *, and one of three nucleosomes in **. (B) Representative histone methyltransferase activity assays with ^14^C-labeled S-adenosylmethionine analyzed by SDS-PAGE, with gels imaged for ^14^C signals (top) or stained with Coomassie blue to confirm equal loading of PRC2 and nucleosomes (bottom). (C) Quantification of three replicates. Error bars are mean ± standard deviation. IC_50_ is the concentration of RNA that inhibits 50% of PRC2 activity relative to no RNA. (D) Table summarizes the four categories of RNA characterized in this study. Affinity (*K_d_*): PRC2-RNA binding affinities measured by EMSA. CLIP in cells: RNA motifs identified by CLIPseq in cells ^30^.

### G4 RNA inhibits PRC2 activity in cells

Unlike the current view in which the ability of RNA to bind PRC2 determines PRC2 inhibition (e.g., poly(A) RNA sequence does not bind PRC2 and therefore does not change PRC2 activity in cells), our biochemical and structural observations defined a group of RNAs that bind PRC2 with similar affinity without inducing PRC2 dimerization or altering PRC2 activity. This led us to propose a new model that in cells, PRC2 constantly engages various RNA sequences, but only some of them can serve as regulators of PRC2 activity.

To test this model, we followed the strategy of Beltran *et al.* ^19^, who validated a CRISPR-dCas9 system to specially position RNA elements in the periphery of the transcription start site of a PRC2-targeted gene, *CDKN2A*, in Malignant Rhabdoid Tumor (MRT) cells (Fig. 5A). In each case, the sgRNA is extended at its 3’ end to display a selected RNA sequence at a genomic locus determined by the CRISPR guide sequence ^55^. We tested five different RNAs (Fig. S12A): sgRNA G4 and sgRNA Poly(A) were the sequences termed G-tract and A-tract in Beltran *et al.*, which contain an array of several G4 clusters or poly(A) clusters, respectively. We designed sgRNA TERRA_mut_ and sgRNA P10 to have multiple repeats of TERRA_mut_ and P10, 220 nucleotides in length (excluding the region of CDKN2A crRNA and tracrRNA), similar to the first two RNAs. Finally, a sgRNA without an extension was included to assess the effect of binding the CRISPR-dCas9 machinery to the gene.

**Figure 5.**
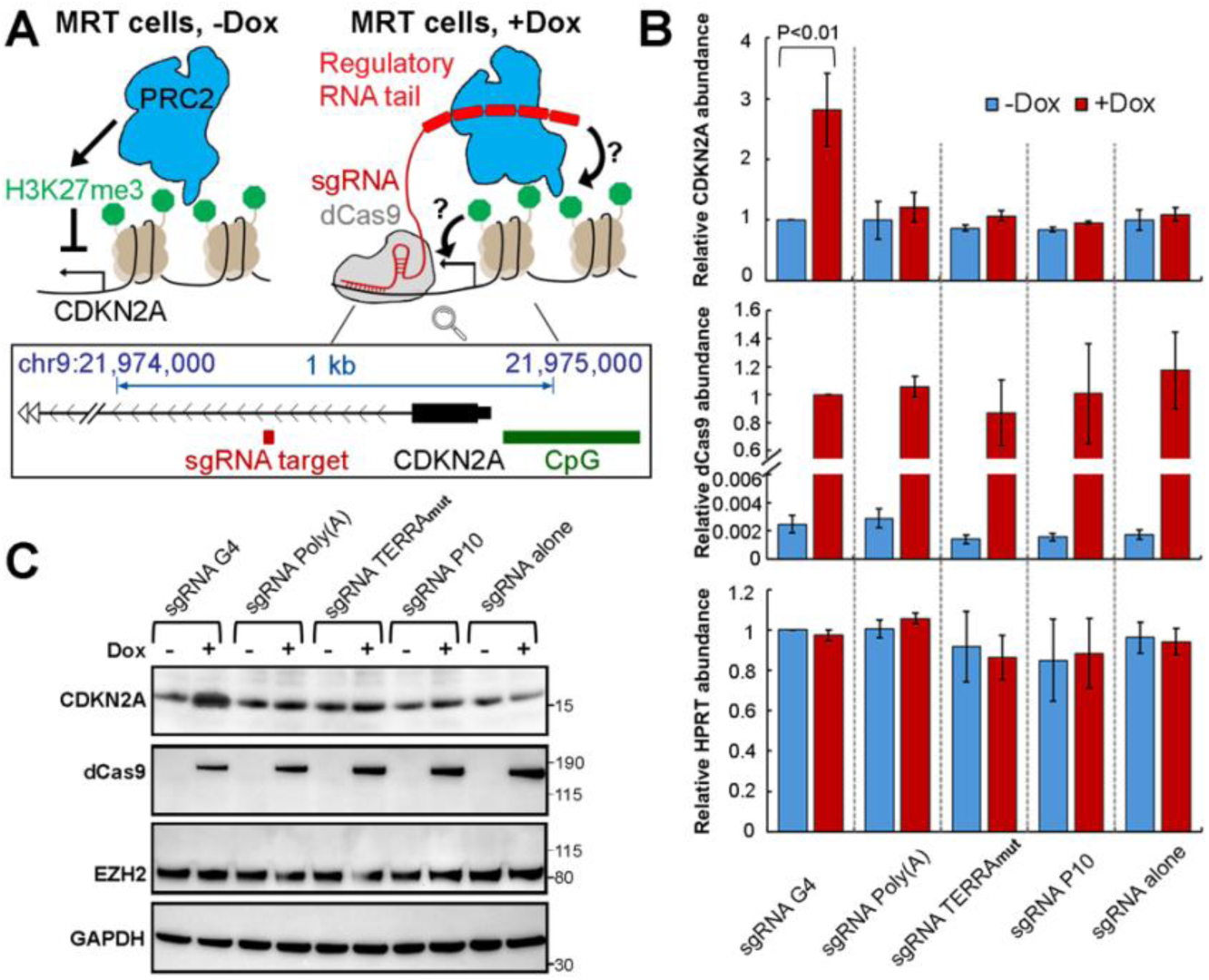
RNA sequence-dependent regulation of PRC2 in cells. (A) Top: schematic of CRISPR-dCas9 strategy to display various RNA “tails” at a specific gene and examine the functional consequences. Bottom: genome browser representation indicates the position of sgRNA target (dark red), *CDKN2A* transcription start site (black) and upstream CpG island (green). (B) RT-qPCR measures abundance of several mRNAs in MRT transgenic cells in the absence or presence of dox. Relative abundance of mRNAs was calculated by ΔΔCq method using GAPDH as internal reference. HPRT is mRNA from another housekeeping gene. Three independent biological replicates were examined, and error bars are mean ± standard deviation. P value was calculated by Student’s t-test. (C) Western blot results of MRT transgenic cells in the absence or presence of dox. Two biological replicates gave equivalent results.

Stable transgenic cells containing a doxycycline (dox)-inducible dCas9 cassette and U6 promoter-driven modified sgRNAs were grown in the presence of dox for 8 days, which led to noticeable production of dCas9 (Fig. 5B and C). Although modified sgRNAs were constitutively expressed, they were stabilized 5-10 fold by the expression of dCas9 (Fig. S12B), consistent with the formation of dCas9 RNPs. Importantly, the repression of *CDKN2A* was relieved when dCas9 and sgRNA G4 were both present (Fig. 5B and C), consistent with G4 RNA being a negative regulator of PRC2 function. None of the other RNAs affected *CDKN2A* gene expression. Although TERRA_mut_ induced PRC2 dimerization and inhibition of activity in vitro, it was inactive in this in vivo system, suggesting that either the lower binding affinity of TERRA_mut_ to PRC2 or the abundance of other competing proteins in the nucleus that bind ssRNA might make TERRA_mut_ insufficient for PRC2 inhibition in this cell system. P10 was selected as an example of a pyrimidine-rich RNA because it was identified as a prevalent PRC2 binding consensus by CLIPseq ^30^. In agreement with our biochemical results, P10 did not mitigate the PRC2-mediated repression of *CDKN2A*. Therefore, PRC2 binds a wide variety of RNA sequences via different binding modes, only some of which result in inhibition, which could contribute to the specificity of PRC2 to only be regulated at certain genes.

### Limitations of UV crosslinking for capturing PRC2-RNA complexes

A recent study failed to capture RNAs bound to PRC2 in living cells ^56^, challenging the widespread reports of PRC2-RNA interaction and, in particular, inconsistent with our model and in vivo results. In this article, Guo *et al.* modified the conventional CLIP assay to a method called CLAP (covalent linkage and affinity purification), which allows denaturing washes before the identification of protein-associated RNA. Like CLIP, CLAP is completely dependent on 254 nm UV crosslinking, whose underlying biophysical and chemical principles are incompletely understood. However, it is unambiguous that protein-RNA photochemical crosslinking is biased towards uracil ^57–59^ and towards certain amino acids ^60–62^, and highly sensitive to the precise distance and orientation of RNA base relative to protein amino acid ^60,61,63,64^. For example, RNA recognition motifs (RRMs), which rely on the stacking interaction between aromatic amino acids and RNA bases, are UV-crosslinked with unusually high efficiency, dominating CLIP-seq databases ^62^. Therefore, we suspected that the efficiency of UV crosslinking might not be sufficient to covalently link PRC2 to G-rich RNAs bound to it in cells.

We therefore performed in vitro UV crosslinking assays of PRC2 simultaneously with the positive control used in Guo *et al.*, namely PTBP1 (Polypyrimidine Tract-Binding Protein 1), which harbors four canonical RRMs and interacts with a defined RNA consensus UCUUUCU ^65^ (Fig. S13). We incubated PTBP1 or PRC2, each with its cognate RNA, at concentrations 10 times greater than the K_d_ values to ensure saturated binding in all reactions. The PTBP1-(UCUUUCU)_6_ interaction was efficiently crosslinked at multiple UV energies, including 0.25 J/cm^2^, which is frequently applied to intact cells and was used in Guo *et al.*. In the PRC2 reactions, we selected the RNA termed 2G4 used in the initial cryo-EM determination of a PRC2-RNA complex, which does not contain any uracil in the RNA sequence. We also examined TERRA and TERRA_mut_ RNAs that contain a few U bases. We found that PRC2 did crosslink to these three RNAs, but with very low efficiency. The RRM-harboring protein PTBP1 crosslinked to associated RNA more than an order-of-magnitude better than PRC2 (Fig. S13D). The intrinsic inefficiency of PRC2-RNA UV-crosslinking may contribute to the failure to capture PRC2-RNA complexes by the CLAP technique.

## DISCUSSION

Several previous studies validate the biological importance of RNA binding in the precise control of PRC2 function in cells ^18–21,39^. Inhibition of PRC2 activity by nascent RNA is thought to help maintain epigenetic gene silencing, ensuring that PRC2 adds its repressive histone mark only at those genes already silenced at the transcriptional level. In contrast, particular lncRNAs have been reported to recruit PRC2 to target genes, serving as positive regulators of PRC2 chromatin occupancy ^13,66^. The sequence and structural diversity of these RNAs could potentially provide the specificity needed by PRC2 reacting differentially, but it also raises the critical question of how PRC2 can recognize different RNAs without a strict sequence consensus.

Here, we show that two categories of RNA can mediate separate binding modes of PRC2, which, importantly, lead to distinct functional consequences. G-rich RNAs, including TERRA_mut_ and pUG-fold, can mediate PRC2 dimerization and inhibit catalytic activity through the same mechanism as canonical G4 RNA, even though the RNA secondary structures are quite different. Our structural analysis indicates that the flexible positively charged loops of PRC2 suspend each of these RNAs between two PRC2 protomers with the ability to accommodate different RNA structures. In contrast, pyrimidine-rich RNAs have strong PRC2 binding affinity without the formation of a PRC2 dimer. This monomeric RNA-binding mode of PRC2 allows PRC2 to simultaneously engage nucleosomes in a manner that does not limit histone H3 tail accessibility or inhibit PRC2 activity. AlphaFold 3 prediction and mutagenesis verification revealed the importance of accessory protein AEBP2 in monomeric PRC2 binding RNA. In summary, this study determines the nucleotide compositions of RNA recognized by PRC2 in two separate binding modes and the molecular basis of RNA sequence-dependent inhibition of PRC2, which reconciles differential regulation of PRC2 by RNA.

Despite G4 RNA being predominantly studied for PRC2-RNA interaction, many independent research groups have investigated other RNA sequences, including HOTAIR lncRNA ^67–69^, Repeat A motif of Xist lncRNA ^32,70^, RNA transcribed from B2 SINE retrotransposon ^71,72^, and artificial sequences without continuous G-tracts ^29,34^. RIPseq and CLIPseq analysis of RNA bound by PRC2 identified C and U repetitive motifs and shorter G-tract motifs incapable of folding into G4 in addition to the G4-forming sequences ^30,40^. Our results support the conclusion that G4 structure is not necessary for RNA to be recognized by PRC2. Multiple RNA sequences bind PRC2 with relatively strong affinity.

Our previous structural characterization of the G4 RNA-bound PRC2 presented the first use of streptavidin-affinity EM grid method for an RNP complex ^22^, a technical advance that selected RNA-bound complexes and enhanced particle quality by holding particles away from the denaturing water-air interface ^41,43^. In this work, we tested this technique with more challenging RNPs composed of less tightly bound RNAs. The successful negative staining and cryo-EM results support the potential of this method to obtain structural information of less preferred minor populations and even transiently interacting complexes. Another new analysis involved integrating AlphaFold3 predictions to illustrate the flexibility of RNA conformations and orientations in our complex. The high similarity between the AlphaFold3 overlapping map and our experimental cryo-EM density supports the accuracy of the predictions and demonstrates how RNA flexibility impacts cryo-EM map quality. Furthermore, it provided clues for testing the role of PRC2 accessory protein AEBP2 in accommodating RNA, which could not be interpreted directly from our cryo-EM map. These new methods are likely to facilitate many future studies of other non-canonical protein-RNA interactions.

Like PRC2, many RNA-binding proteins (RBPs) utilize intrinsically disordered arginine- or lysine-rich patches to engage various RNAs with limited sequence specificity (see reviews ^73–75^). In more than 20% of RBPs, intrinsically disordered regions have been found to comprise a significant portion of the protein sequence. Therefore, it is essential to recognize that the identification of multiple RNA motifs for a given RBP does not necessarily imply that all resulting RNPs perform similar functions. Although conventional methods, including CLIP-seq ^76^ and RIP-seq ^26^ are powerful tools for mapping RNA-protein interactions, they cannot resolve the functional heterogeneity among distinct RNA motifs. As demonstrated in this study, the functional outcomes of RNA binding to PRC2 can depend strongly on the sequence and structural properties of the RNA itself. This emphasizes the importance of detailed biochemical and structural analyses to determine and elucidate how specific RNA sequences modulate the activity or behavior of their RBP partners.

## LIMITATIONS OF THE STUDY

Although the streptavidin-affinity EM method has notable benefits of selecting desired particles and preventing damage by the water-air interface, it has the intrinsic disadvantage of applying multiple layers of support background, significantly decreasing signal intensity. We found that it had a preferred application on large-sized particles (i.e., dimeric PRC2) and negative-stained particles but not small-sized particles (i.e., monomeric PRC2). Second, although we used specialized cryo-EM preparation and processing methods, TERRA_mut_ RNA and peripheral protein regions in our density map did not reach high resolution. We cannot unambiguously model TERRA_mut_ and demonstrate the exact amino acid residues within PRC2 flexible loops responsible for RNA binding. In addition, AlphaFold3 could not predict RNA interactions with PRC2 subunits with high confidence (chain iPTM and chain-pair iPTM < 0.2). Although the predicted AEBP2-RNA interactions are consistent between models, the details of this interaction differ. Our data cannot provide an atomic model to illustrate how zinc-finger domains of AEBP2 bind RNA. Furthermore, PRC2 holoenzyme assembles with various accessory proteins to form multiple subcomplexes in cells (see review ^77^). The recombinant PRC2 used for in vitro assays mainly focused on a particular PRC2.2 complex, which contains an embryo-specific isoform of AEBP2 and a truncated JARID2. In our modified CRISPR-dCas9 experiment, we do not know which specific subcomplex(es) are responsible for the repression of *CDKN2A* in MRT cells.

Finally, our results do not address the possibility that in some cases of sparse transcription, RNA might be able to recruit PRC2 to chromatin (reviewed in ^13,66^), an activity that is biophysically possible ^78^ but difficult to test in cells.

## ACKNOWLEDGEMENTS

We thank E. Hartwick and S. Laursen (University of Colorado Boulder Biochemistry Krios Electron Microscopy Facility RRID: SCR_019057) for cryo-EM data collection and storage infrastructure maintenance. We thank S. Zimmermann and G. Morgan (University of Colorado Boulder Electron Microscopy Service) for negative staining data collection. We thank A. Erbse (University of Colorado Boulder Shared Instruments Pool RRID SCR_018986) for providing research instrumentation, in particular the Avanti JXN 26 centrifuge (funded by NIH Grant R24OD033699-01). We thank T. Nahreini (University of Colorado Boulder Cell Culture Facility RRID:SCR_018988) for use of the cell culture facility.

## DATA AND MATERIALS AVAILABILITY

Cryo-EM density maps and fitted models have been deposited in the Electron Microscopy Data Bank (EMD-46751, consensus map; EMD-46722, Body1 from multibody refinement; and EMD-46726, Body2 from multibody refinement) and the Protein Data Bank (PDB: 9DCH). Requests for reagents, plasmids, cell lines, and zebrafish strains used in this study should be directed to the corresponding authors.

## STAR★METHODS

### KEY RESOURCES TABLE

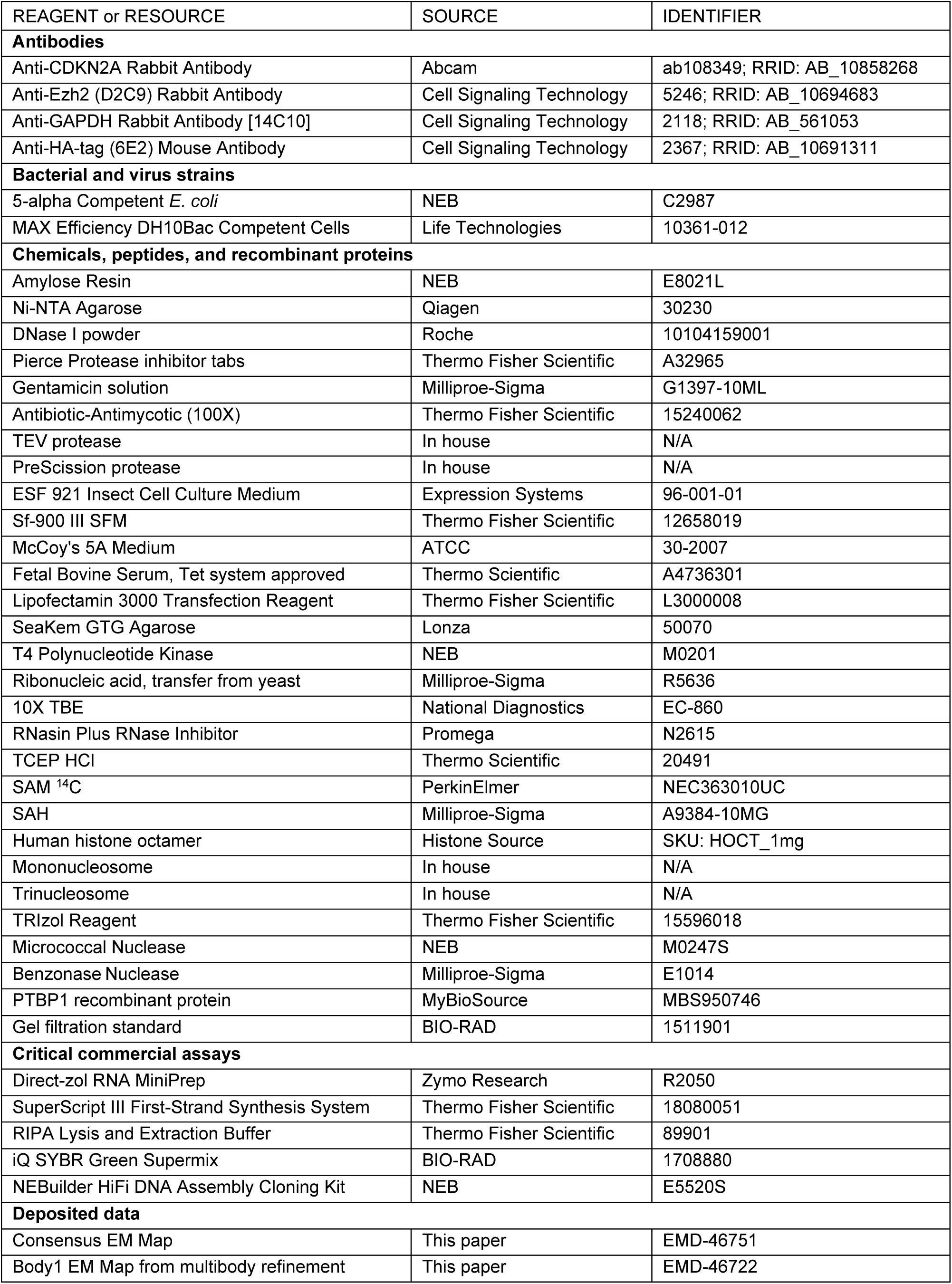
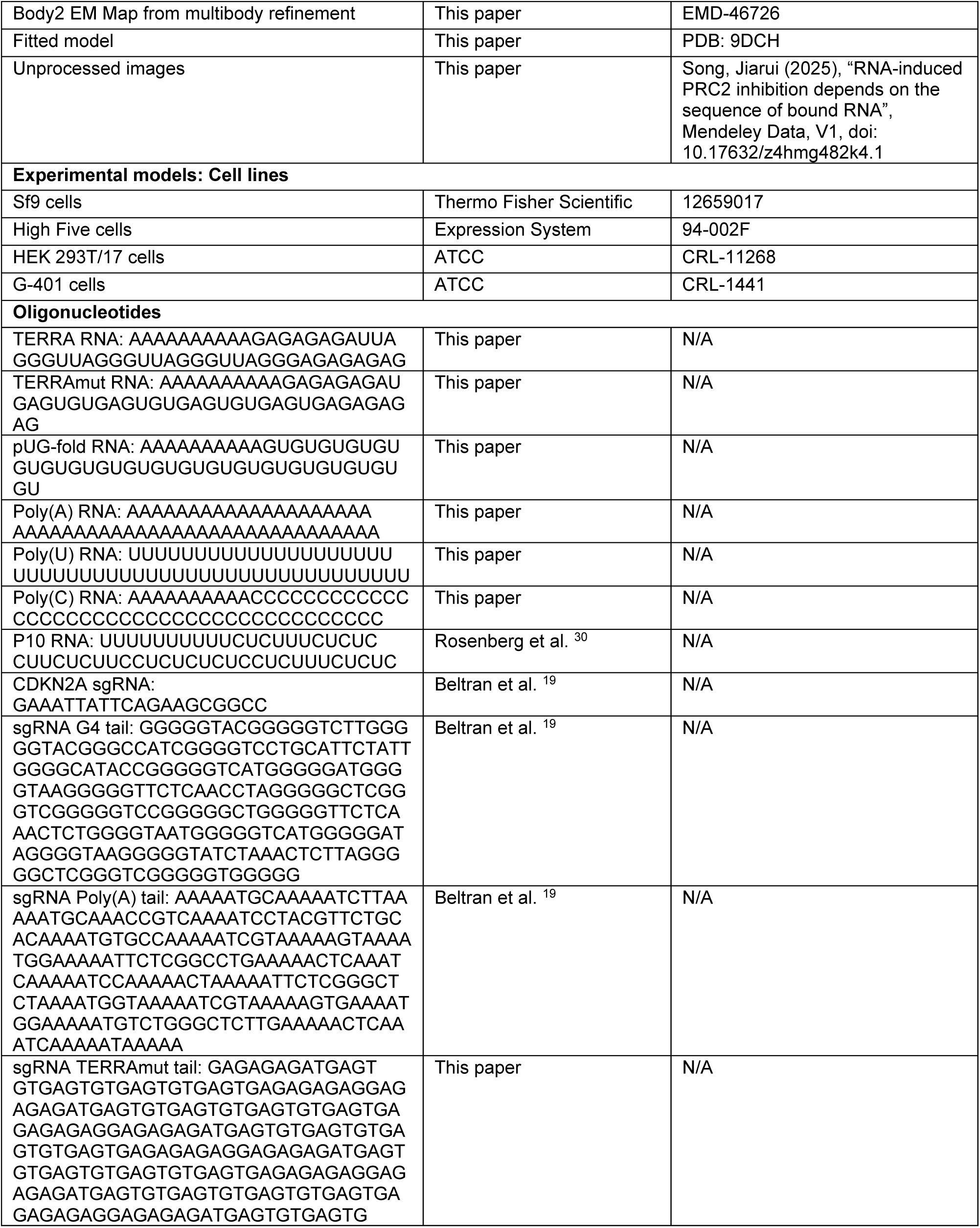
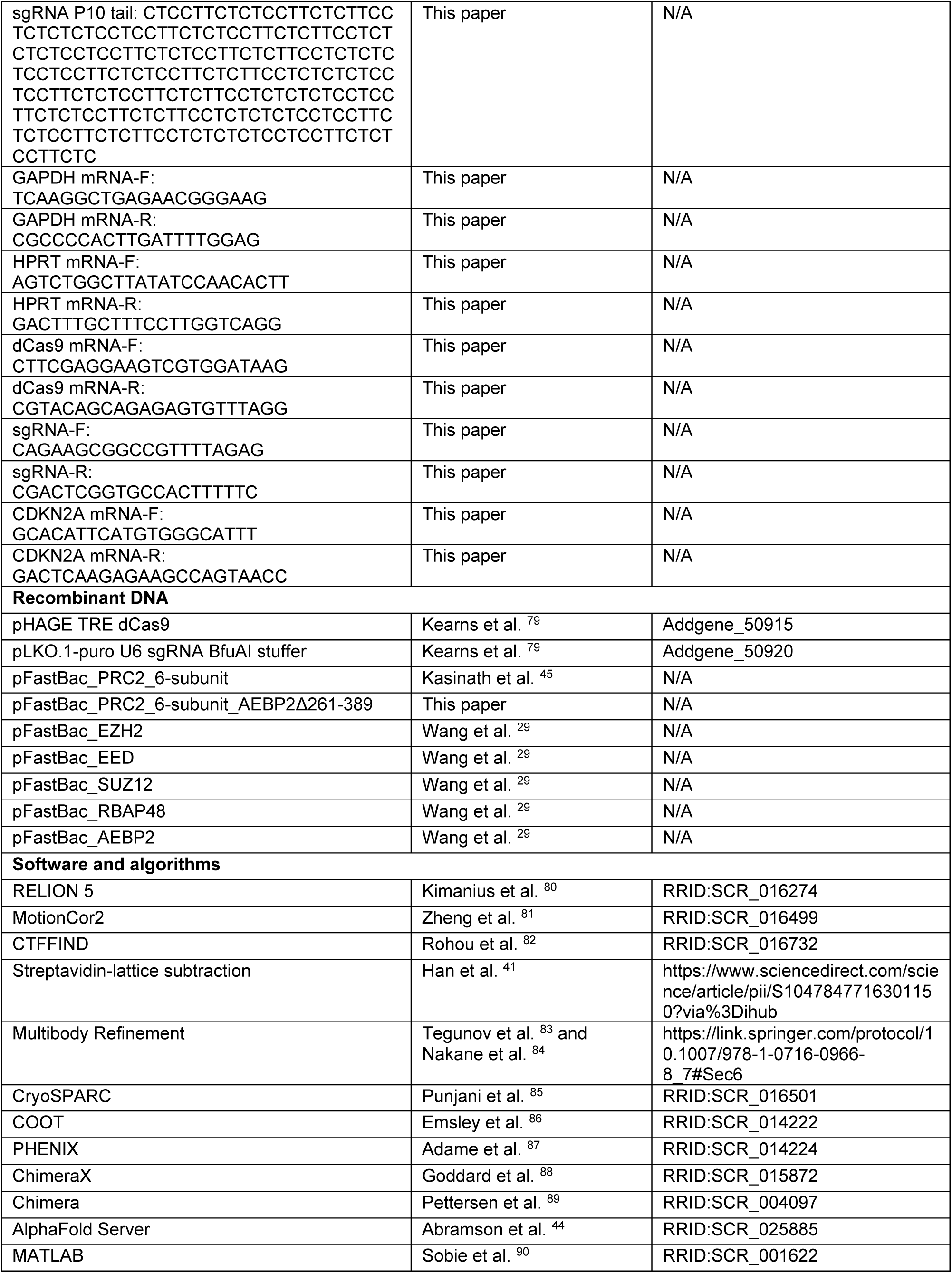
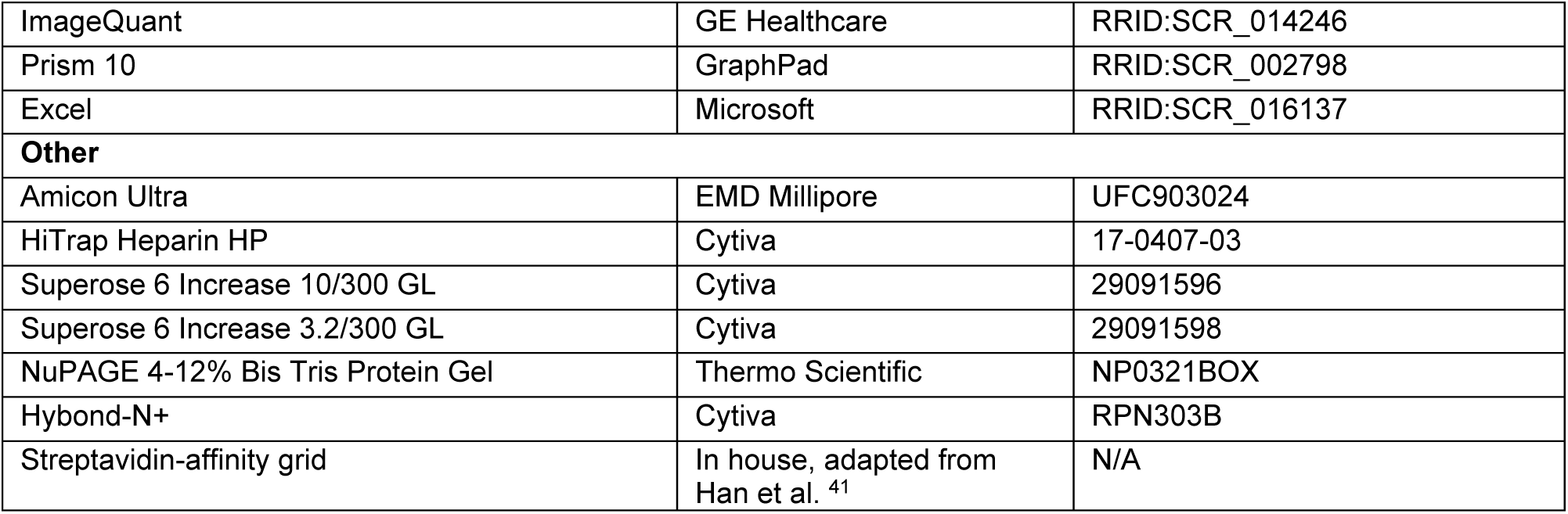

### EXPERIMENTAL MODEL AND STUDY PARTICIPANT DETAILS

#### HEK 293T/17 cell culture

HEK 293T/17 cells were obtained from and authenticated by ATCC (CRL-11268, LOT: 63696280) and maintained in Dulbecco’s Modified Eagle Medium (DMEM) supplemented with 10% fetal bovine serum (FBS), 1X GlutaMAX, pyruvate, and penicillin/streptomycin at 37 degree and 5% CO_2_. Cells were routinely tested for mycoplasma contamination by the University of Colorado Boulder Cell Culture Facility (RRID:SCR_018988).

#### G-401 cell culture

Malignant Rhabdoid Tumor cells (G-401) were obtained from and authenticated by ATCC (CRL-1441, LOT: 70055408) and maintained in McCoy’s 5A media (ATCC, 30-2007) supplemented with 10% Tet-approved FBS (Thermo Scientific, A4736301) and penicillin/streptomycin at 37 degree and 5% CO_2_.

### METHOD DETAILS

#### Protein expression and purification

For the preparation of the six-subunit PRC2, genes encoding full-length EED, SUZ12, RBAP48, His-tagged EZH2 isoform 2 (UniProt Q15910-2), Strep-GFP-tagged embryonic isoform of AEBP2, and Strep-GFP-tagged truncated JARID2 (amino acids 119-450) were cloned into a single multi-bac plasmid ^45^. Each expression cassette had an independent promoter and terminator. This multi-bac plasmid was used to make infectious baculovirus stock in Sf9 (*Spodoptera frugiperda*, IPLB-Sf-21-AE) cells using the Bac-to-Bac system (Invitrogen). Then, HighFive (*Trichoplusia ni*, BTI-Tn-5B1-4) cells were transfected with baculovirus at 28°C for 66 hours to express recombinant complex. Cells were washed with cold PBS buffer and frozen in liquid nitrogen until use.

All purification steps were performed in a 4°C cold room. Cells were lysed in lysis buffer (25 mM HEPES pH 7.9 at 4°C, 250 mM NaCl, 2 mM MgCl_2_, 1 mM TCEP, 10 mM imidazole, 0.5% NP-40, 10% glycerol, DNase I and protease inhibitor cocktail) for 1 hour and sonicated. Debris was then removed by centrifugation at 15,000 rpm for 35 min. The supernatant was incubated with Ni-NTA agarose resin (Qiagen) for 1 hour, and resin was washed with 10 column volumes (CV) of lysis buffer, 10 CV of high-salt wash buffer (25 mM HEPES pH 7.9 at 4°C, 1 M NaCl, 2 mM MgCl_2_, 1 mM TCEP, 0.01% NP-40, and 10% glycerol), and 20 CV of low-salt wash buffer (25 mM HEPES pH 7.9 at 4°C, 150 mM NaCl, 2 mM MgCl_2_, 1 mM TCEP, 30 mM imidazole, and 10% glycerol). Proteins were then eluted in elution buffer (25 mM HEPES pH 7.9 at 4°C, 150 mM NaCl, 2 mM MgCl_2_, 1 mM TCEP, 300 mM imidazole, and 10% glycerol) and dialyzed twice (1 hour each) in buffer (25 mM HEPES pH 7.9 at 4°C, 150 mM NaCl, 2 mM MgCl_2_, 1 mM TCEP, and 10% glycerol) to remove imidazole. Proteins were incubated with TEV protease overnight after concentrating to 3-5 mg/ml. The AKTA-FPLC system was used for subsequent purification with a HiTrap Heparin HP column (Cytiva) and a Superose 6 increase 10/300 column (GE Healthcare). Heparin column was equilibrated with buffer I (20 mM HEPES pH 7.9 at 4°C, 150 mM NaCl, 2 mM MgCl_2_, 1 mM TCEP, and 10% glycerol), and sample was eluted with a linear gradient of buffer II (20 mM HEPES pH 7.9 at 4°C, 2 M NaCl, 2 mM MgCl_2_, 1 mM TCEP, and 10% glycerol). The Superose 6 increase 10/300 column was equilibrated and run with final storage buffer (25 mM HEPES pH 7.9 at 4°C, 150 mM KCl, 2 mM MgCl_2_, 10% glycerol, and 1 mM TCEP). Protein complex was flash frozen in liquid nitrogen as single-use aliquots and stored at −80°C.

Preparations of five-subunit PRC2 and four-subunit PRC2 were similar to six-subunit PRC2 with several modifications. Four-subunit PRC2 contained full-length EED, SUZ12, RBAP48, and EZH2 that were all MBP-tagged. Five-subunit PRC2 had the subunits included in the four-subunit with addition of MBP-tagged short isoform of AEBP2.

Amylose agarose resin was used to replace the Ni-NTA resin in the initial affinity purification of the complexes. After washing with lysis buffer (same lysis buffer without imidazole), high-salt wash buffer, and low-salt wash buffer (same buffer without imidazole), PRC2 was eluted with elution buffer (low-salt wash buffer with 10 mM maltose). Without dialysis, the eluate was directly concentrated and then digested by Prescission protease at 4°C overnight to remove MBP tags. Further purification involving the HiTrap Heparin HP column and Superose 6 increase 10/300 column was identical to six-subunit PRC2. The protein complex was flash-frozen in liquid nitrogen as single-use aliquots and stored at −80°C.

#### RNP complex assembly

All RNA oligos were purchased from Dharmacon Custom RNA Synthesis (Horizon), including HPLC purification service. TERRA, TERRA_mut_, pUG-fold, Poly(A), Poly(U), Poly(C) and P10 RNAs were synthesized in two versions. RNA oligos without modification had 5’-hydroxyl ends for ^32^P radiolabeling, while 5’-biotinylated RNAs were used for structural studies compatible with streptavidin-affinity EM grids. To promote RNA folding, all RNAs including single-stranded RNAs were heated at 95°C for 2-3 min, snap-cooled on ice for 5 min, then refolded in RNP complex buffer (25 mM HEPES pH 7.9 at 4°C, 50 mM KCl, 2 mM MgCl_2_, 10% glycerol, and 1 mM TCEP) at 37°C for 20 min. PRC2 and S-adenosylhomocysteine (SAH) were added into the reaction at final concentrations of 600 nM and 40 µM, respectively, and the reaction was incubated at 30°C for 30 min to assemble the RNP complex.

#### EM sample preparation

Quantifoil Au 1.2/1.3 grids were converted to streptavidin-affinity grids in-house using procedures previously described ^41,43^. Grids were re-hydrated in EM preparation buffer I (25 mM HEPES pH 7.9 at 4°C, 50 mM KCl, 2.5% glycerol, and 1 mM TCEP) at room temperature (RT) for 1 hour. After removing the remaining buffer, 4 µl of the assembled RNP complex was applied to the surface of the streptavidin-affinity grid. The grid was incubated for 5-10 min in a humidified chamber, washed with 40 µl of EM preparation buffer I, and then washed with 40 µl of EM preparation buffer II (25 mM HEPES pH 7.9 at 4°C, 50 mM KCl, 2.5% glycerol, 0.01% NP-40, and 1 mM TCEP). After the washes, the buffer was wicked away using Whatman filter paper, and 4 µl of the EM preparation buffer II was added immediately. The grid was then transferred to the Leica EM GP2 plunge freezer, blotted for 2-3 s at 10°C and 90% humidity, and then plunged into liquid ethane. Negative staining of the streptavidin-affinity grid followed the same protocol, but instead of a plunge freezer, five droplets of 40 µl uranyl formate (30 mg/mL) stain were used.

#### EM data collection and processing

Cryo-EM data were collected using a Titan Krios G3i equipped with a Thermo Fisher Falcon 4 direct-electron detector (DED) camera and a Selectris energy filter set with a 10-eV slit width. Data acquisition was performed using Thermo Fisher EPU at 130,000x magnification (0.97 Å/pixel) with a defocus range of −1.9 to −0.5 µm. Movies were collected in EER format with a total dose of 50 electrons per square angstrom (e^−^/Å^2^) and an exposure time of 5.49 s corresponding to 1323 frames. Gain correction was applied during motion correction using Relion’s own implementation of the UCSF motioncor2 program. The same parameters were used for ± 20° tilted stage data collection.

Negative staining datasets were collected on a Tecnai F20 microscope operated at 200 kV, with a Gatan K3 direct detector, at a nominal magnification of 25,000x, corresponding to 1.449 Å per pixel. Datasets were collected using a dose of 40-60 e^−^/Å^2^ on streptavidin-affinity grids with 3 nm carbon supports on the back of the grids.

Data were processed in RELION 5 ^80^. The movie frames were aligned using RELION’s own (CPU-based) implementation of the UCSF MotionCor2 program ^81^ and CTF parameters were fit using CTFFIND ^82^. The background streptavidin lattice of each micrograph was subtracted using in-house scripts ^41^. TOPAZ automatic picking was trained on selected particles and applied to pick all micrographs. Initial models were generated within RELION from negative staining data and used as reference for the first round of 3D classification. Later classifications used references from previous good classes. Subsequent processing steps included several runs of regular 3D classification with Blush regularization, particle subtraction built in RELION 5, and 3D classification without alignment (regularization parameter T=24) (Figure S4). The selected 120,658 particles were then re-extracted and subjected to per-particle defocus refinement, beam-tilt refinement, and 3D refinement with Blush regularization to generate the consensus map. Soft-edged masks of individual PRC2 protomers were applied in the multibody refinement ^83,84^ to improve map qualities. Local resolution estimation was performed in RELION 5 using the same soft, spherical masks used during refinement. Local resolution filtered maps were generated in CryoSPARC ^85^.

#### Model building

Individual PRC2 protomers were built using cryo-EM maps from the multibody refinement. The coordinates of G-quadruplex RNA-bound PRC2 six-subunit complex (PDB: 8FYH) provided a starting model from which all the coordinates were adjusted and rebuilt in the new map using COOT ^86^. The model of each PRC2 promoter was subjected to global refinement and minimization in real space using PHENIX ^87^. These were then subjected to manual inspection and adjustment in COOT, followed by refinement again in PHENIX. TERRA_mut_ RNA model was generated using a 10-nucleotide single-stranded fragment (UGAGUGUGAG) from AlphaFold3 prediction model 1 and then docked into our map for the position we designated as the RNA density. The cryo-EM density maps and the molecular graphics were prepared with Chimera and ChimeraX ^89^.

#### Generating representative EM map from AlphaFold3 predicted models

PRC2 subunit sequences and TERRA_mut_ RNA sequence were provided to the AlphaFold Server for structure prediction. The top four AlphaFold3 models were modified in Chimera by ‘molmap’ command to generate corresponding maps with an 8 Å low-pass filter. All four maps were added to a combined map by the ‘vop add’ command. In parallel, ‘Segger’ function was used to create surface models with one segmentation region for each individual map. Then, the combined map was subtracted four times, and only density within every surface model was kept. Therefore, the final map retained overlapping regions without flexible areas that differed between predictions.

#### Electrophoretic mobility shift assay (EMSA)

RNA oligos were radiolabeled at 37°C for 30 min using T4 polynucleotide kinase (NEB, M0201L). After labeling, the oligos were purified on a denaturing polyacrylamide gel. The bands were excised from the gel and eluted with TE, then precipitated with NaCl, glycogen, and EtOH. Pellets were resuspended in TE, and the counts of the oligos were determined by liquid scintillation counting. Radiolabeled oligos were heated, snap-cooled, and refolded in EMSA binding buffer (50 mM Tris-HCl, pH 7.5 at 25°C, 100 mM KCl, 2.5 mM MgCl_2_, 0.1 mM ZnCl_2_, 2 mM 2-mercaptoethanol, 0.05 mg/ml BSA, 0.05 mg/mL yeast tRNA, and 5% glycerol). Next, stock PRC2 was diluted with EMSA binding buffer to a series of concentrations and mixed with the radiolabeled oligos. Binding was carried out at 30°C for 30 min, and then samples were loaded onto a non-denaturing 1.0% agarose gel (Lonza SeaKem GTG agarose) buffered with 1X TBE. Gel electrophoresis was for 90 min at 66 V in a 4°C cold room. A Hybond N^+^ membrane (Amersham, Fisher Scientific 45-000-927) and two sheets of Whatman 3 mm chromatography paper were put underneath the gel, then the assembled gel was vacuum dried for 60 min at 80°C. Dried gels were exposed to phosphorimaging plates and scanned using a Typhoon phosphorimager (GE Healthcare) for signal acquisition. Gel analysis was carried out with ImageQuant software (GE Healthcare), and data were fitted to the nonlinear binding curve (specific binding with Hill slope) using Prism software.

#### Native gel electrophoresis

8% polyacrylamide (29:1 ratio of acrylamide to bisacrylamide), 0.5X TBE and 100 mM KCl were mixed before pouring into a 0.75 mm thickness gel cassette. RNA samples were radiolabeled and refolded as described above. After mixing with loading dye (0.5X TBE, 10% glycerol, and a trace amount of bromophenol blue and xylene cyanol), 1000 cpm of each RNA sample was loaded. Native gel was run at room temperature in running buffer (0.5X TBE and 100 mM KCl) at 50-60 volts to prevent any substantial increase in temperature. After electrophoresis, two sheets of Whatman 3-mm chromatography paper were put underneath the gel, and then the assembled gel was vacuum-dried for 45 min at 80°C. Dried gels were exposed to phosphorimaging plates and scanned using a Typhoon phosphorimager (GE Healthcare) for signal acquisition.

#### Analytic size-exclusion chromatography

In a 50 μL reaction, 2 μM PRC2 (final concentration) and 2 μM refolded RNA (final concentration) were mixed with RNP complex buffer (25 mM HEPES pH 7.9 at 4°C, 50 mM KCl, 2 mM MgCl_2_, 10% glycerol, and 1 mM TCEP). The reaction was incubated at 30°C for 30 min to complete RNP assembly and then injected into a Superose 6 increase 3.2/300 column (Cytiva) pre-equilibrated with the RNP complex buffer. The column was run at a flow rate of 0.02 ml/min, monitored by UV260 and UV280 detectors. Gel filtration standard (Bio-rad, #1511901) was injected and run using the same protocol to estimate the molecular weight of monomeric and dimeric PRC2 complexes.

#### Fluorescence resonance energy transfer (FRET)

40 µL reactions were prepared in FRET Binding Buffer (50 mM Tris-Cl pH 7.5 at 25°C, 100 mM KCl, 2.5 mM MgCl_2_, 0.1 mM ZnCl_2_, 0.1 mg/mL nonacetylated BSA, 5% glycerol, 2 mM 2-mercaptoethanol), with 200 nM each of Cy3-labeled TERRA and Cy5-labeled Poly(C) RNAs and 0-2.4 µM PRC2 protein. Control reactions were also performed with individual RNAs and with no ligand to rule out FRET-independent fluorescence effects. All reactions were incubated for 1 h at room temperature in a black, 384-well microplate (Corning, #3575), then analyzed with a TECAN Spark microplate reader. Fluorescence properties measured included Cy3 fluorescence and anisotropy (Ex = 512 ± 30 nm, Em = 567 ± 20 nm), Cy5 fluorescence and anisotropy (Ex = 620 ± 30 nm, Em = 675 ± 20 nm), and FRET (Ex = 512 ± 30 nm, Em = 675 ± 20 nm).

Anisotropy binding curve data were fit with Equation 1, where *A* is anisotropy, [*E_T_*] is total PRC2 concentration, *n* is Hill coefficient, and *K_D_* is PRC2-ligand equilibrium dissociation constant.

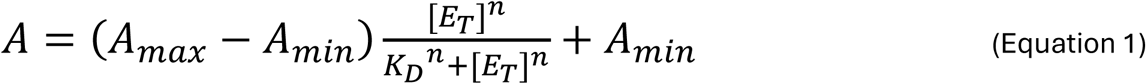

FRET behavior under an independent binding model was predicted from binding curve data via Equation 2, where *F* is relative FRET signal, [*L_T_*] is ligand concentration (each, assuming Cy3- and Cy5-labeled ligands are equal), [*E_T_*] is protein concentration, *Kd_1_* and *n_1_* are the Equation 1 coefficients from the Cy3-labeled ligand binding curve data, and *Kd_2_*and *n_2_* are the Equation 1 coefficients from the Cy5-labeled ligand binding curve data.

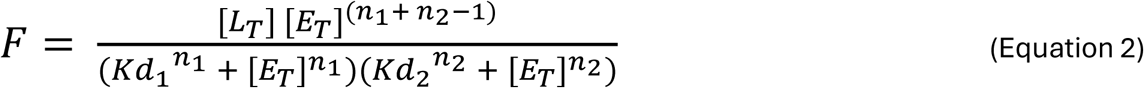

#### RNA-competition EMSA

RNA oligos were ^32^P-radiolabeled as described in the conventional EMSA method. Radiolabeled oligos were heated, snap-cooled, and refolded in EMSA binding buffer (50 mM Tris-HCl, pH 7.5 at 25°C, 100 mM KCl, 2.5 mM MgCl_2_, 0.1 mM ZnCl_2_, 2 mM 2-mercaptoethanol, 0.05 mg/ml BSA, 0.05 mg/mL yeast tRNA, and 5% glycerol). PRC2 (0.5 µM final concentration) was added and incubated at 30°C for 20 min to achieve RNP assembly. Then, non-radiolabeled competitor RNA was refolded and diluted with EMSA binding buffer to corresponding concentrations and mixed with the previous reactions. Competition was carried out at 30°C for 30 min, and then samples were loaded onto a non-denaturing 1.0% agarose gel (Lonza SeaKem GTG agarose) buffered with 1X TBE. Gel electrophoresis, gel drying process and signal acquisition were the same as described for conventional EMSA.

#### Nucleosome-RNA competition assay

We prepared ^32^P-radiolabeled trinucleosomes in-house by adding a small fraction of radiolabeled DNA into non-labeled DNA during trinucleosome assembly. DNA is 621 bp in total, including a 33 bp 5’ tail, three 143 bp Widom sequences, two 64 bp linkers between Widom sequences, and a 31 bp 3’ tail. Unmodified Human octamer was purchased from The Histone Source (SKU: HOCT). For nucleosome-RNA competition, 600 nM PRC2, 100 nM labeled trinucleosome, and serial dilutions of refolded RNA were combined in RNP complex buffer (25 mM HEPES pH 7.9 at 4°C, 50 mM KCl, 2 mM MgCl_2_, 10% glycerol, and 1 mM TCEP) and incubated at 30°C for 30 min. In this assay, unlike RNA-competition EMSA, PRC2 was not pre-incubated with labeled nucleosomes. All three components in the reactions were combined simultaneously. After incubation, samples were loaded onto 1% agarose gel (SeaKem GTG Agarose) buffered with 1X TBE and resolved at 66 V for 110 min in a 4°C cold room. Gels were vacuum dried for 60 min at 80°C. Dried gels were exposed to phosphorimaging plates and signal acquisition was performed using a Typhoon Trio phosphorimager (GE Healthcare).

#### Methyltransferase activity assay

Mononucleosomes were prepared in house by assembling 192 bp DNA (21 bp tail + 143 bp Widom sequence + 28 bp tail) and unmodified Human octamer (Histone Source, SKU: HOCT). 600 nM PRC2 and serial dilutions of refolded RNAs were pre-incubated at 30°C for 20 min to reach binding equilibrium and then assembled into methyltransferase reaction mix including 1X methyltransferase buffer (25 mM HEPES pH 7.9 at 4°C, 50 mM KCl, 2 mM MgCl_2_, 10% glycerol, and 1 mM TCEP), 0.1 mg/ml BSA, 1X protease inhibitor, 1 µl RNase inhibitor/20 µl solution, 10 µM ^14^C SAM (PerkinElmer), and 300 nM mononucleosome. The volume of each reaction was 20 µl. Methyltransferase reactions were incubated at 30°C for 20 min prior to boiling with SDS loading buffer to inactivate PRC2. Proteins were separated through NuPAGE 4-12% gel (Invitrogen) by running at 180 V for 52 min. The gel was vacuum dried at 80°C for 30 min and then exposed to phosphorimaging plates. Signal intensities were quantified by ImageQuant, and IC_50_ values were calculated by equation (log(inhibitor) vs. response -- variable slop) built in the Prism software.

#### UV crosslinking assay

PTBP1-binding RNA and PRC2-binding RNAs were radiolabeled as described above. All RNAs were heated to 95°C for 2-3 min, snap-cooled on ice for 5 min, diluted to 20,000 cpm with RNP complex buffer (25 mM HEPES pH 7.9 at 4°C, 50 mM KCl, 2 mM MgCl_2_, 10% glycerol, and 1 mM TCEP), and refolded at 37°C for 20 min. Then, 600 nM PTBP1 or 600 nM PRC2 was added to reactions with corresponding RNAs at a final volume of 110 µl and incubated at 30°C for 30 min to reach binding equilibrium. Then, 20 µl droplets of each reaction were loaded onto a glass cover slide on top of a cooling metal in an ice basket, exposed directly inside a UV crosslinker (UVP, CL1000) with various crosslinking energies. The four different reactions with the same crosslinking energy were exposed simultaneously and then transferred to tubes containing an SDS loading buffer. The glass cover slide was replaced with a new one prior to the next round of crosslinking to avoid sample contamination. Proteins were separated through a NuPAGE 4-12% gel (Invitrogen) by running at 150 V for 60 min. Gel was vacuum dried at 80°C for 30 min, and then exposed to phosphorimaging plates. Signal intensities were quantified by ImageQuant and plotted by Microsoft Excel.

#### Modified CRISPR-dCas9 assay

Malignant Rhabdoid Tumor (MRT) cells (G-401) were obtained from ATCC (CRL-1441) and maintained in McCoy’s 5A media (ATCC, 30-2007). This commercially available media has been modified to contain 1.5 mM L-glutamine and 2200 mg/L sodium bicarbonate. Therefore, only 10% FBS and penicillin/streptomycin were supplemented. Lentivirus was generated in HEK293 cells using third generation packaging and envelope plasmids (Addgene) with pHAGE TRE dCas9 transfer plasmid (Addgene plasmid #50915, a gift from R. Maehr and S. Wolfe, University of Massachusetts Medical School, ^79^). MRT cells were transfected and selected with 200 μg/ml G418 (Geneticin) for 10 days, then maintained in 100 μg/ml G418 for two weeks to amplify enough dCas9 stable cells for further transfections of sgRNA constructs. sgRNA constructs were generated in two steps. First, plasmid synthesis service (GenScript) synthesized initial pUC57 plasmids containing U6 promoter, CDKN2A crRNA sequence, tracrRNA sequence, various RNA sequence extensions (see key resource table for complete sequences), T6 terminator, and multiple restriction enzyme recognition sites including an EcoRI site at the end. Second, the pUC57 plasmids were digested by NdeI (recognition site within U6 promoter sequence) and EcoRI and ligated into pLKO.1-puro U6 sgRNA BfuAI stuffer transfer plasmid (Addgene plasmid #50920, a gift from R.

Maehr and S. Wolfe, University of Massachusetts Medical School, ^79^). Lentivirus generation was the same as before. MRT cells containing dCas9 were transfected and selected with 1 μg/ml puromycin for 10 days. Then cells recovered for 4 days in media without selection prior to doxycycline (Dox) induction. dCas9 expression was induced by 2 μg/mL Dox for 8 days. Media were changed every two days and cells were passed once when reaching high confluency. Total RNA was extracted using Direct-zol RNA miniprep kit (Zymo Research). Total protein was extracted by incubating cells with RIPA lysis and extraction buffer (Thermo Fisher) supplemented with Pierce Protease Inhibitor (Thermo Fisher), 1 mM CaCl_2_, and Micrococcal Nuclease at room temperature for 20 min.

### QUANTIFICATION AND STATISTICAL ANALYSIS

All statistical analyses and definition of replicates are described in the corresponding figure legends. Statistical tests were performed by Student’s T test using GraphPad T test calculator.

