## Supplemental figure S1-S13 and supplemental table S1 for "RNA-induced PRC2 inhibition depends on the sequence of bound RNA"

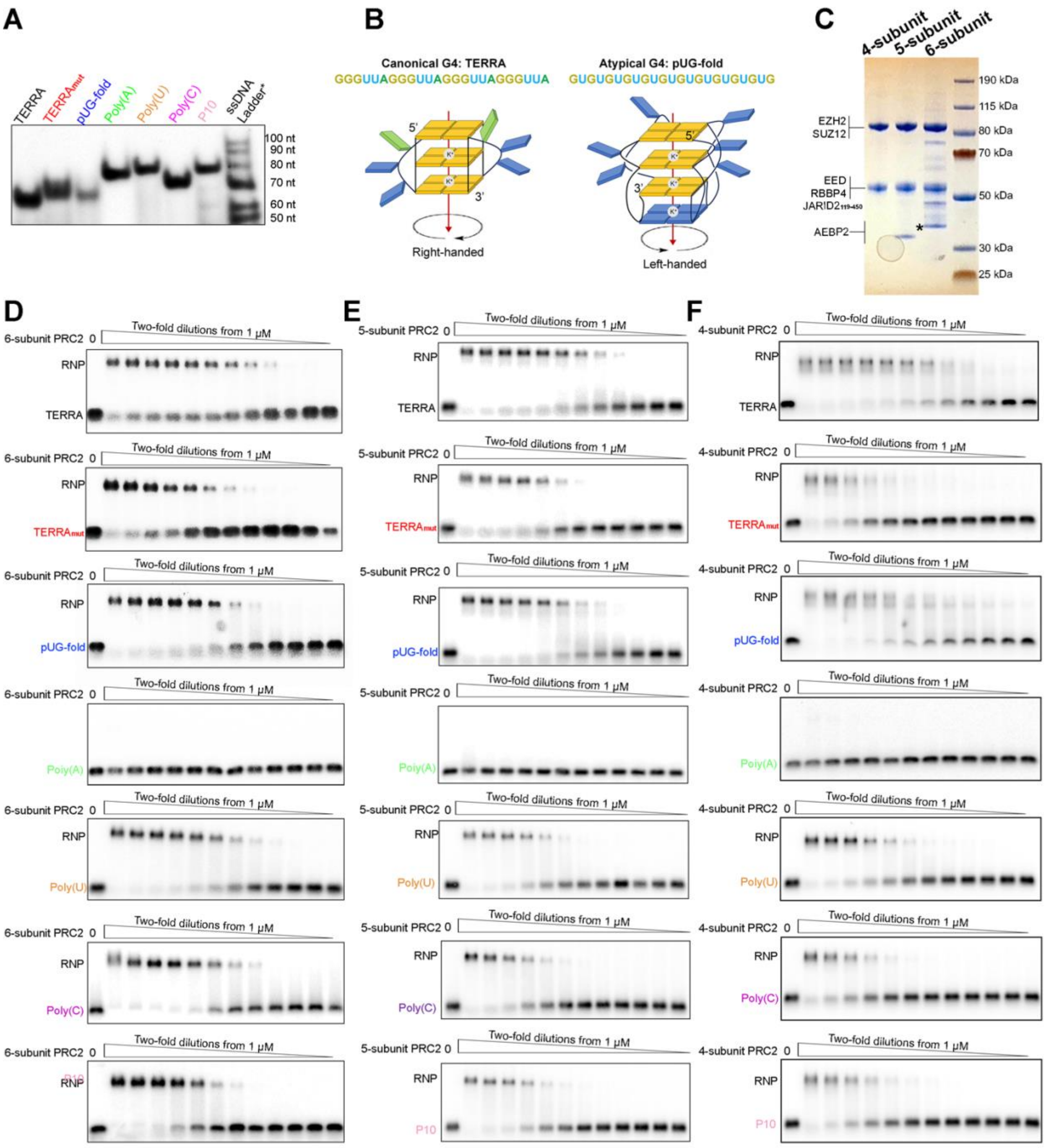


**Figure S1. Six-, five- and four-subunit PRC2 complexes bind RNAs of various sequences and structures.** (A) Native gel electrophoresis of RNA. RNAs were folded in $K^+$ buffer and then analyzed using a native polyacrylamide gel run in $K^+$-containing buffer. *The size marker in this assay was single-stranded DNA ladder (IDT, 20/100 Ladder) which is not an accurate size-reference for RNAs, only serving as a reference for reproducibility. (B) Schematic representations of canonical G4 (TERRA) and atypical G4 (pUG-fold), adapted from [1]. (C) Coomassie-stained gel of purified four-, five-, and six-subunit PRC2 complexes. *AEBP2 of the six-subunit PRC2 has seven additional amino acids at N-terminal after tag cleavage, which is unstructured flexible linker without changing AEBP2 function. (D-F) Representative EMSA gels of RNAs binding to the PRC2 complexes.

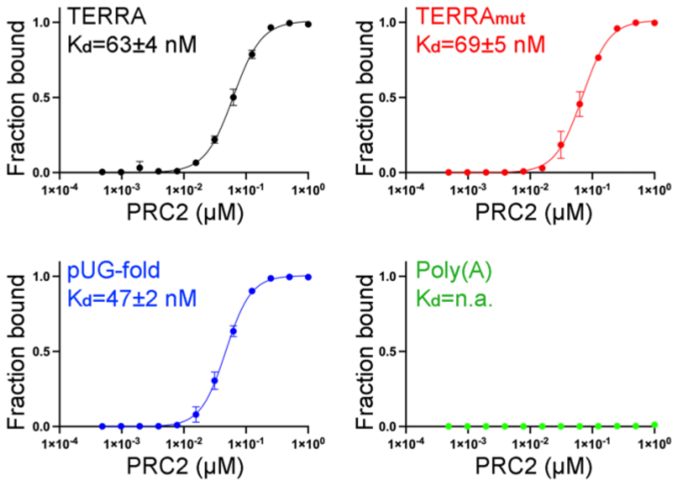


**Figure S2. G-quadruplex formation contributes to the higher binding affinity of TERRA to PRC2.** Quantification of EMSA results in a $Li^+$ reaction buffer in which the G-quadruplexes are not folded. The binding affinity of TERRA is reduced to the same affinity as single-stranded $TERRA_{mut}$ RNA. Error bars are range of two replicates.

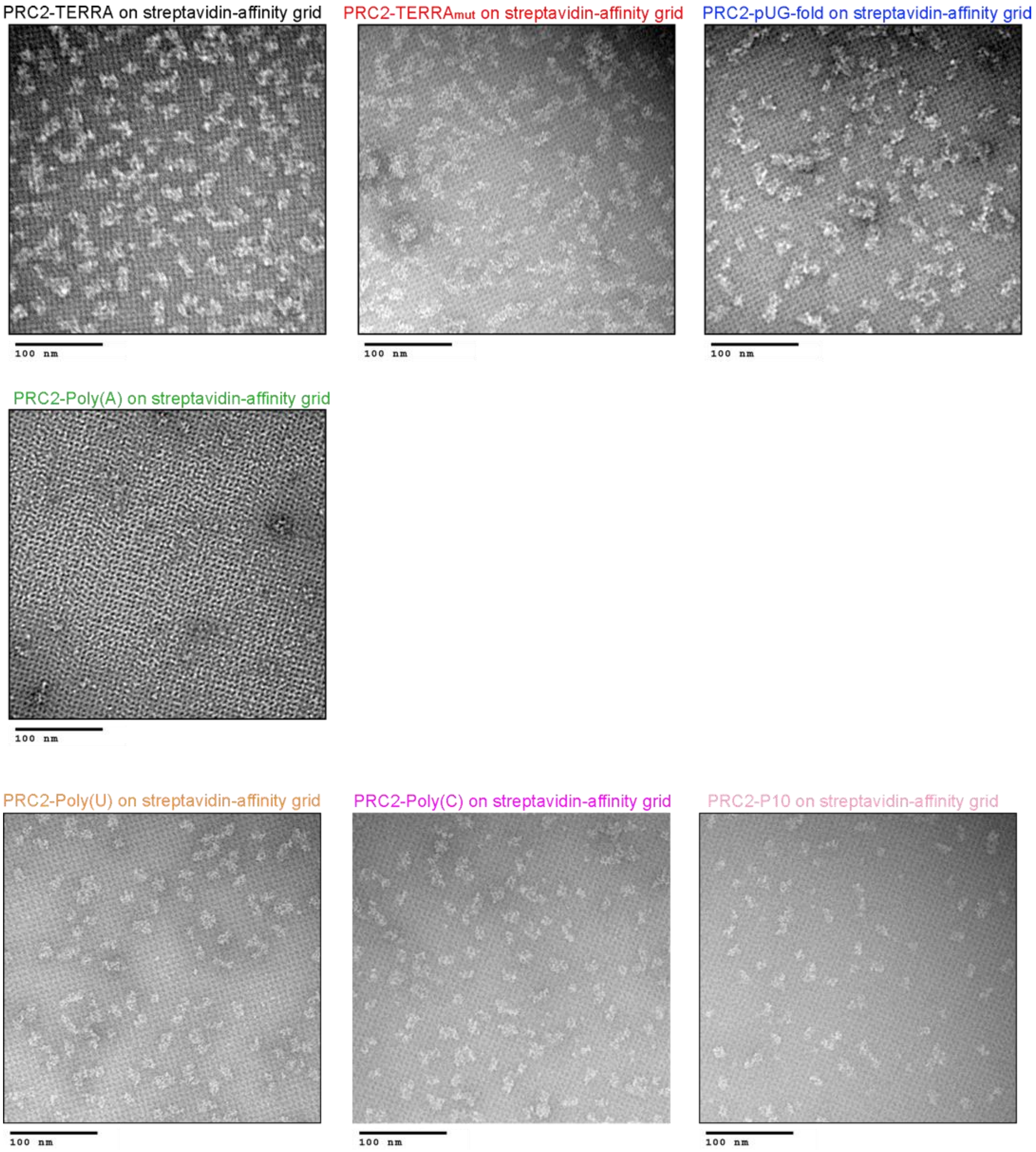


**Figure S3. Streptavidin-affinity EM method selects for RNA-bound PRC2 six-subunit complex.** Representative negative staining EM images of PRC2-TERRA, PRC2-TERRA$_{mut}$, PRC2-pUG-fold, PRC2-Poly(A), PRC2-Poly(U), PRC2-Poly(C), and PRC2-P10. All RNA are biotinylated. Poly(A) does not bind PRC2, and therefore, few recognizable particles are observed in the image.

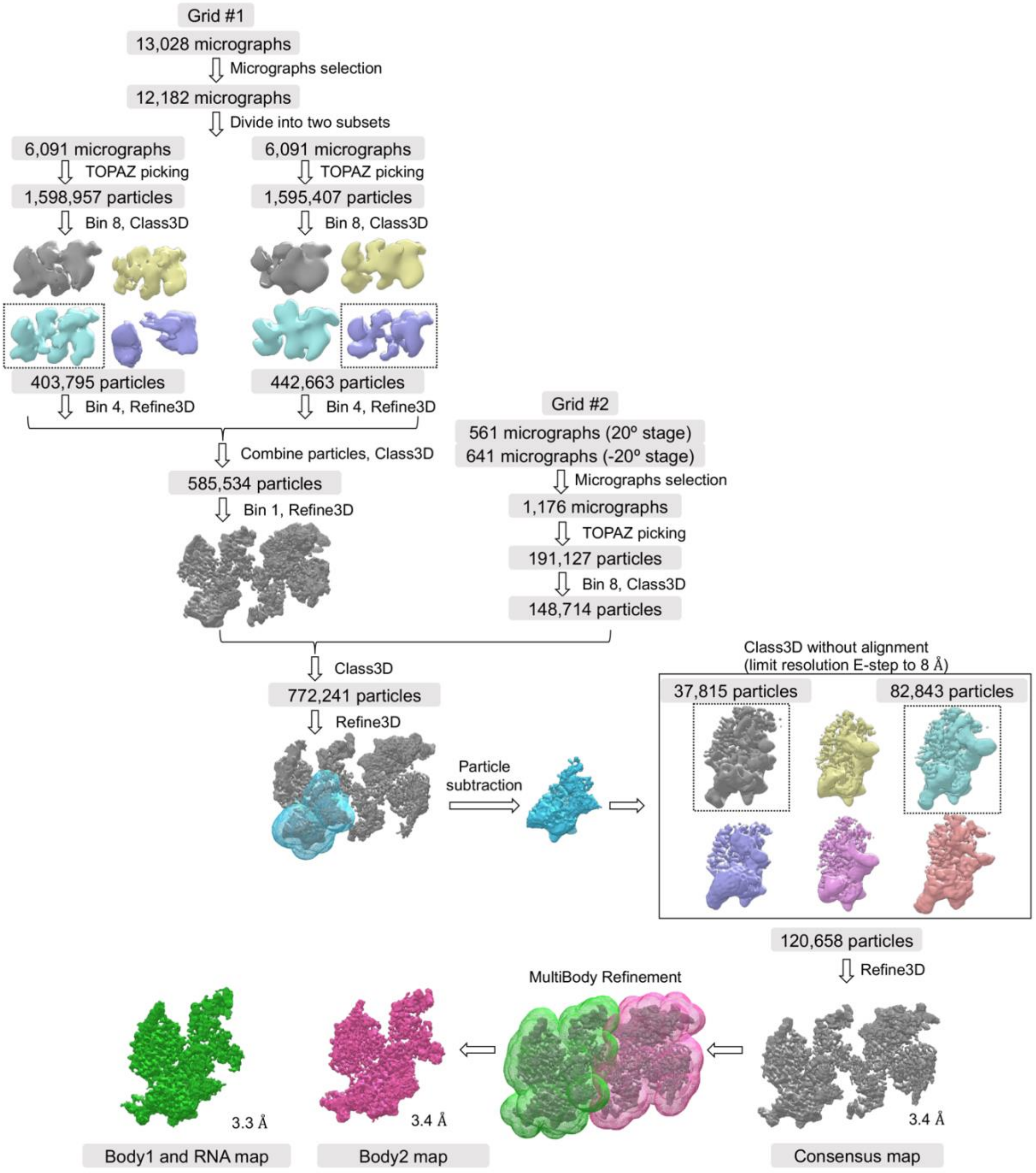


**Figure S4. Single-particle cryo-EM image processing workflows for PRC2-TERRA$_{mut}$ RNA complex.**

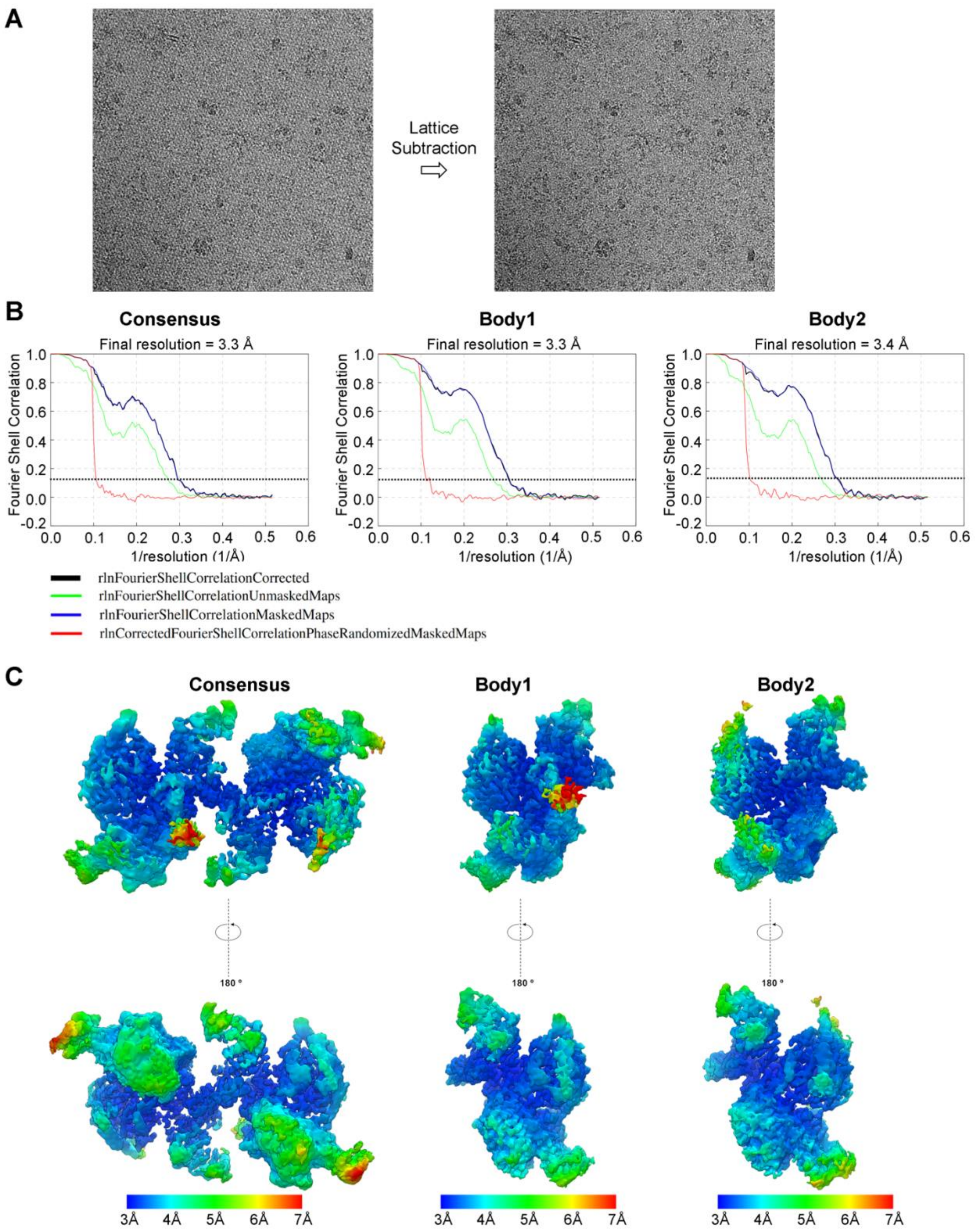
A
Lattice Subtraction
B
Consensus
Final resolution = 3.3 Å
Body1
Final resolution = 3.3 Å
Body2
Final resolution = 3.4 Å
Fourier Shell Correlation
1/resolution (1/Å)
rlnFourierShellCorrelationCorrected
rlnFourierShellCorrelationUnmaskedMaps
rlnFourierShellCorrelationMaskedMaps
rlnCorrectedFourierShellCorrelationPhaseRandomizedMaskedMaps
C
Consensus
Body1
Body2
180 °
3Å 4Å 5Å 6Å 7Å

D

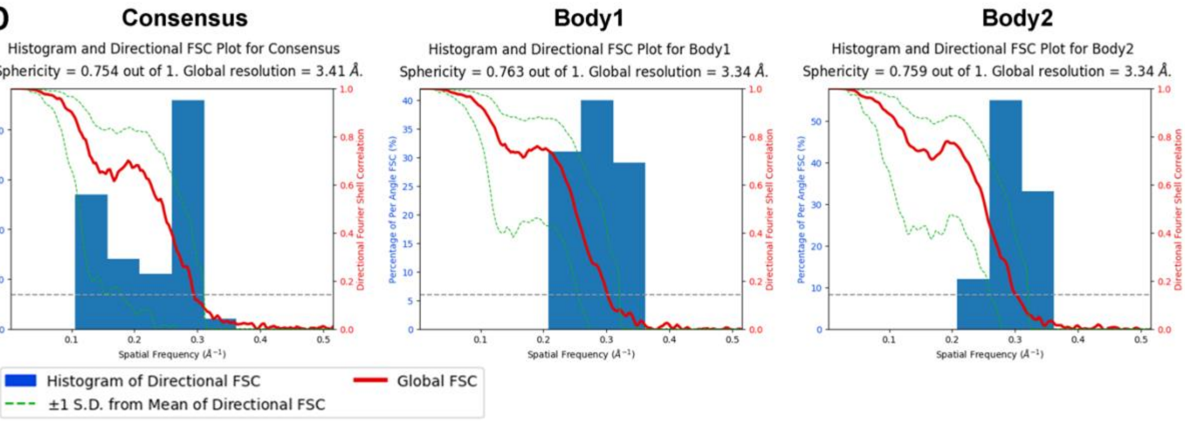

Consensus
Histogram and Directional FSC Plot for Consensus
Sphericity = 0.754 out of 1. Global resolution = 3.41 Å.
Body1
Histogram and Directional FSC Plot for Body1
Sphericity = 0.763 out of 1. Global resolution = 3.34 Å.
Body2
Histogram and Directional FSC Plot for Body2
Sphericity = 0.759 out of 1. Global resolution = 3.34 Å.
Percentage of Per Angle FSC (%)
Directional Fourier Shell Correlation
Spatial Frequency (Å⁻¹)
Histogram of Directional FSC
Global FSC
±1 S.D. from Mean of Directional FSC


E

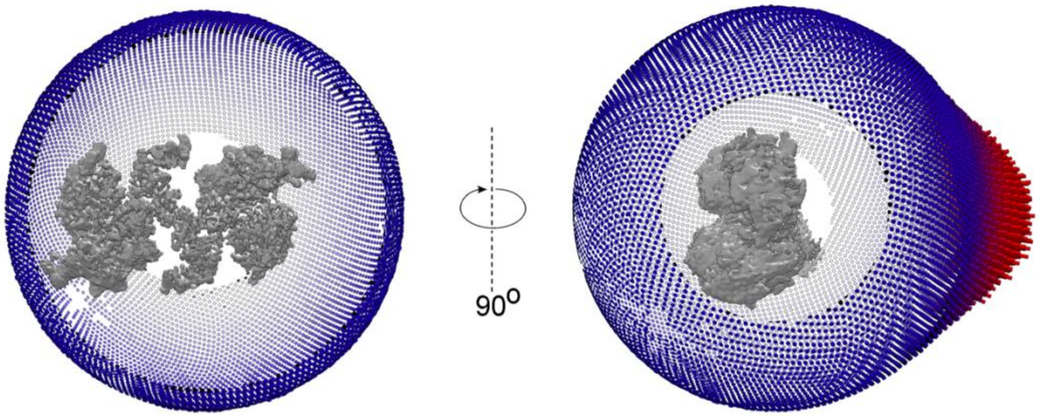

90o


F

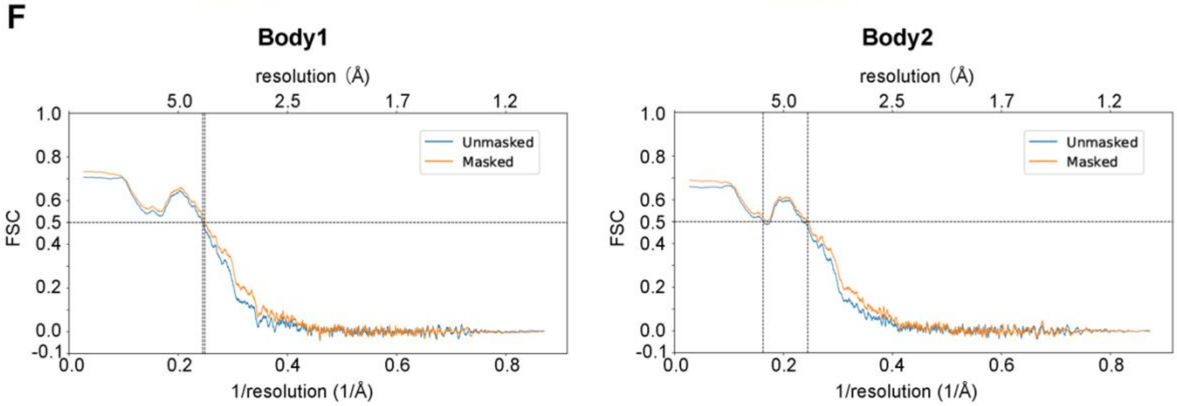

Body1
Body2
resolution (Å)
5.0
2.5
1.7
1.2
FSC
Unmasked
Masked
1/resolution (1/Å)


G

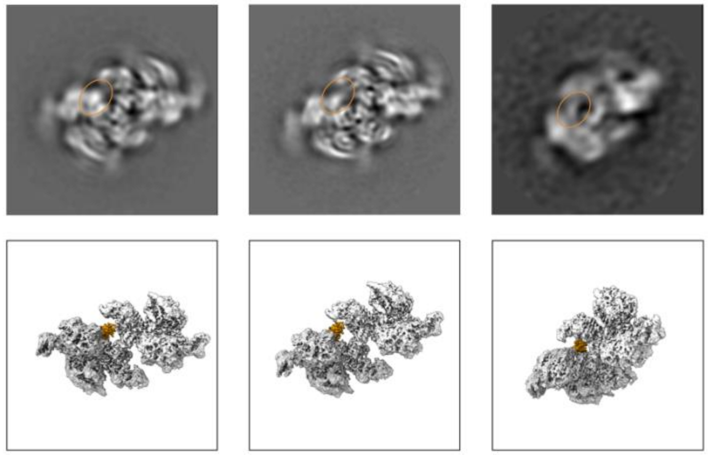

**Figure S5. Cryo-EM map analysis.** (A) Cryo-EM micrographs before and after lattice subtraction. (B) Fourier shell correlation (FSC) curves of consensus map and maps of two individual bodies from multibody refinement. 0.143 intercepts are indicated by dashed lines. (C) Local-resolution density maps of consensus and two bodies. (D) 3D FSC of consensus map and two bodies. (E) Euler angle distribution for the particles after consensus map refinement. (F) Model vs Map FSC for Phenix-refined models of two bodies. (G) Top: A subset of 2D class averages from the final particles used to generate the consensus map. Bottom: 3D density maps at the corresponding orientations. $TERRA_{mut}$ RNA density is highlighted by orange circles in 2D averages and orange color in 3D maps, respectively.

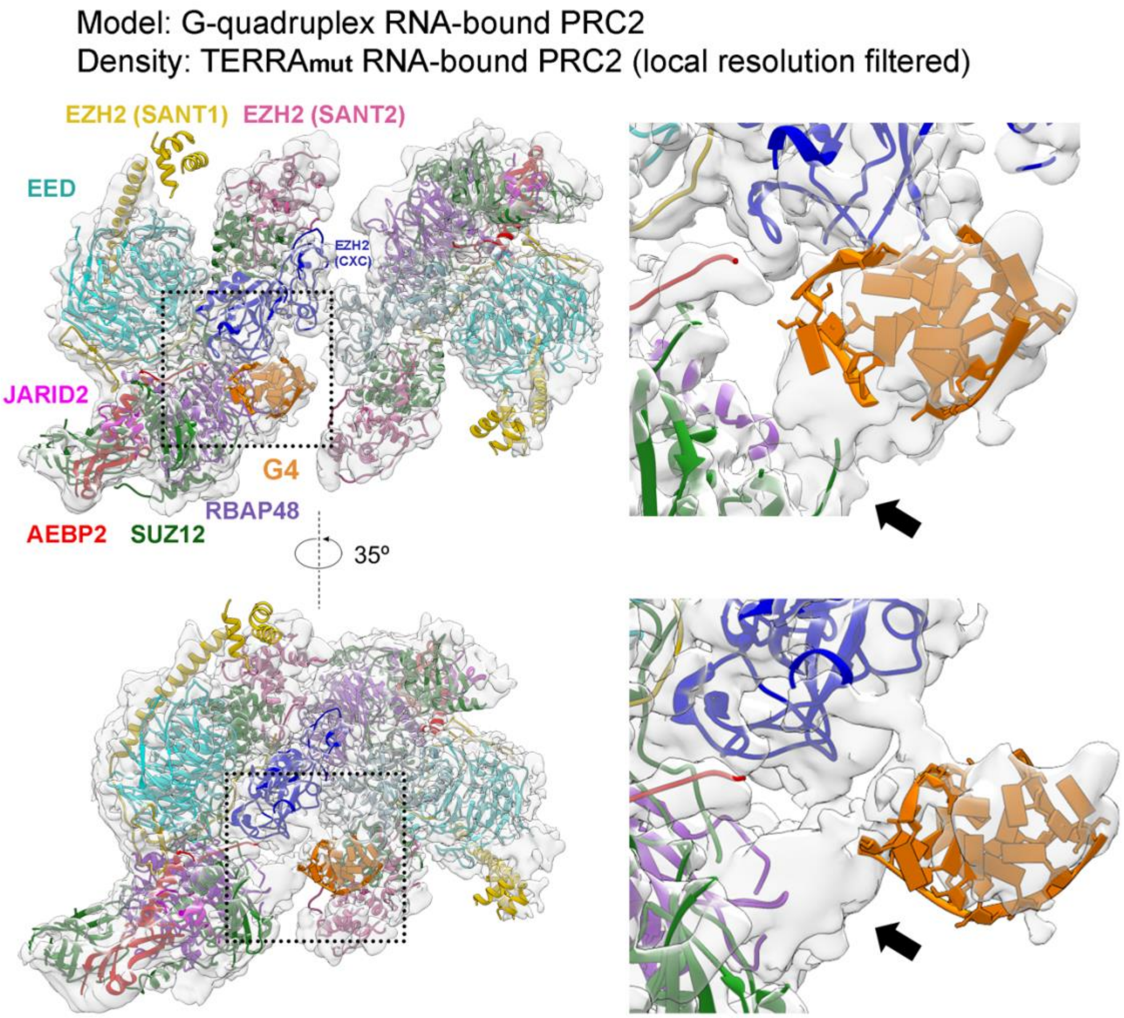


**Figure S6. Direct comparison of G-quadruplex RNA- and $TERRA_{mut}$ RNA-bound PRC2 dimers.** Model is from G-quadruplex RNA-PRC2 complex (PDB:8fyh). Transparent density is the cryo-EM map of $TERRA_{mut}$ RNA-PRC2 complex, local resolution filtered. Dashed boxes are zoomed in to present details at right. Subunits of dimerized PRC2 are at the same arrangement in both structures, while $TERRA_{mut}$-PRC2 complex has the RNA density partially overlapping with G-quadruplex model and additional density of RBAP48 or AEBP2 visualized (arrow).

### AlphaFold 3 predictions of PRC2-TERRAmut RNA

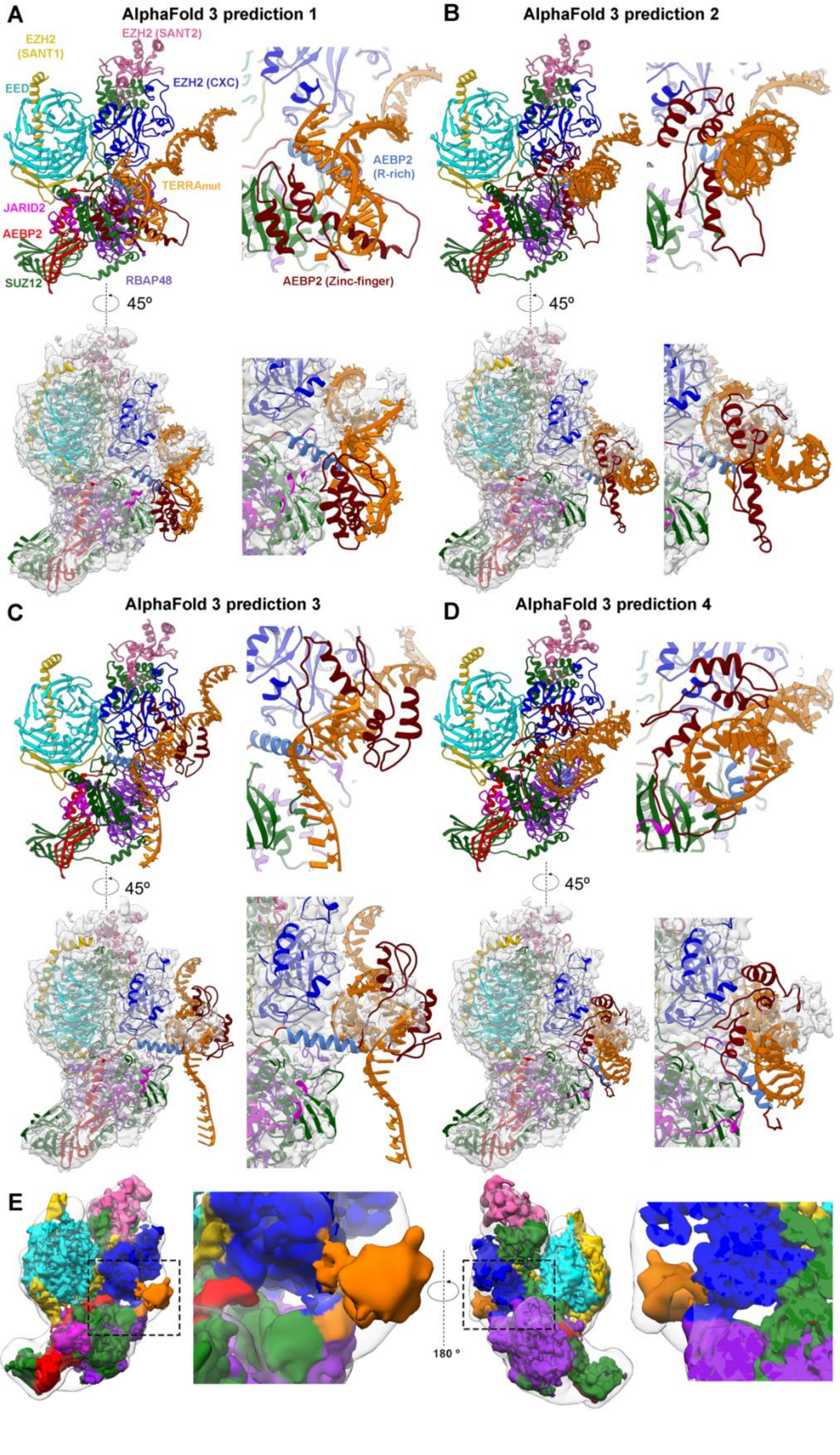

**Figure S7. AlphaFold3 generated predictions of the $TERRA_{mut}$ RNA-single PRC2 complex.** (A) Top left: Model from AlphaFold3 prediction. Top right: Close-up view of $TERRA_{mut}$ RNA and peripheral protein regions. Bottom left: Model is overlapped onto the cryo-EM density to emphasize the intrinsic flexibility of PRC2-$TERRA_{mut}$ binding. Bottom right: Zoom-in view. (B-D) Three additional AlphaFold3 predictions displayed as in panel A. (E) Local resolution filtered cryo-EM map of $TERRA_{mut}$ RNA-bound PRC2 protomer (colored) is overlapped to the AlphaFold3 map (transparent). Dashed boxes are zoomed in to present details.

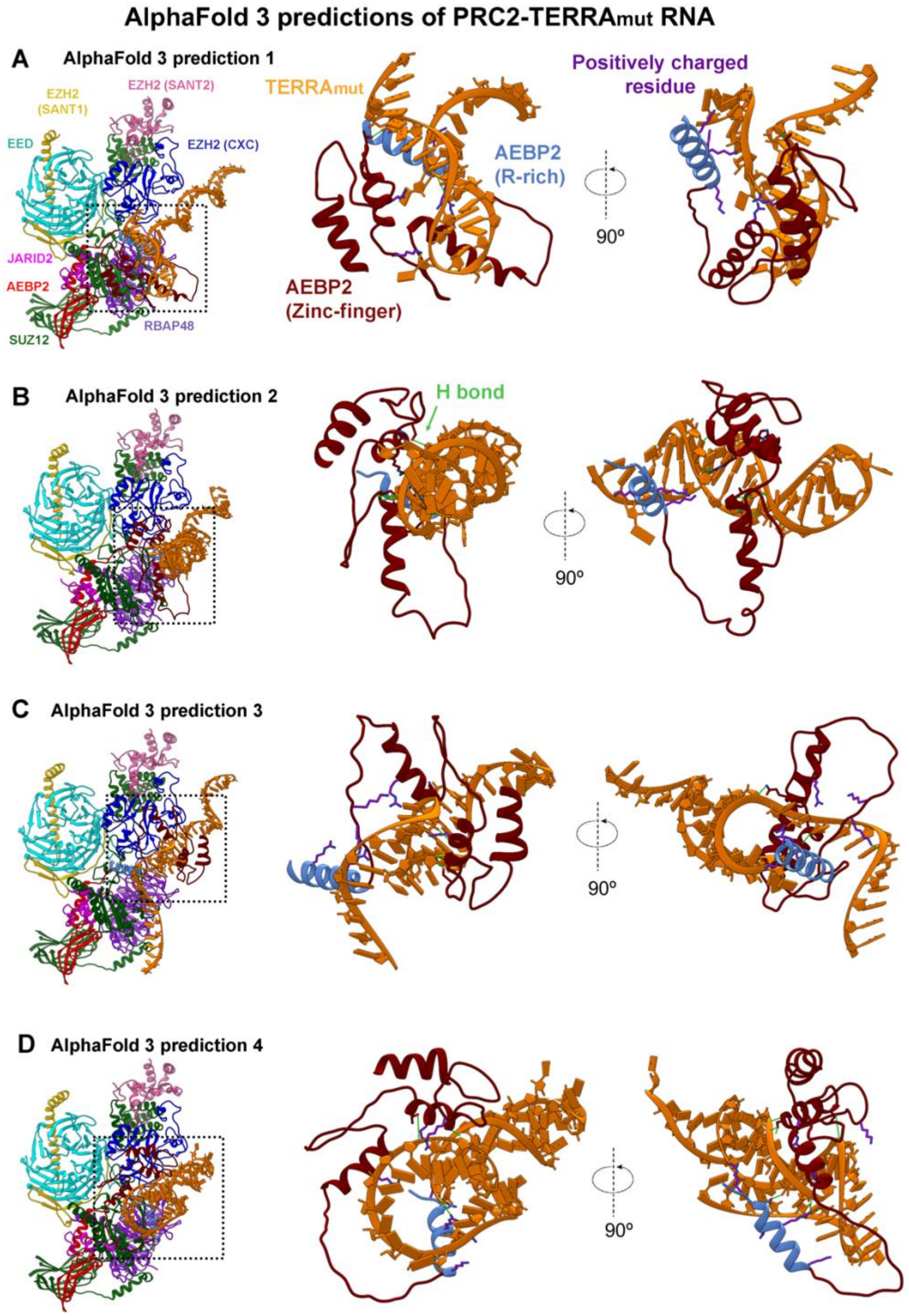


**Figure S8. AlphaFold 3 predicted interactions between AEBP2 and TERRA$_{mut}$ RNA.** (A-D) Left: Overall models from AlphaFold 3 predictions. Dashed boxes are zoomed in. Center and Right: Zoom-in views of TERRA$_{mut}$ RNA, AEBP2 arginine-rich segment, and zinc-finger domains. Hydrogen bonds were analyzed by the FindHbond function built in Chimera and highlighted in green color. Side chains of positively charged residues aiming towards RNA are also highlighted in purple color.

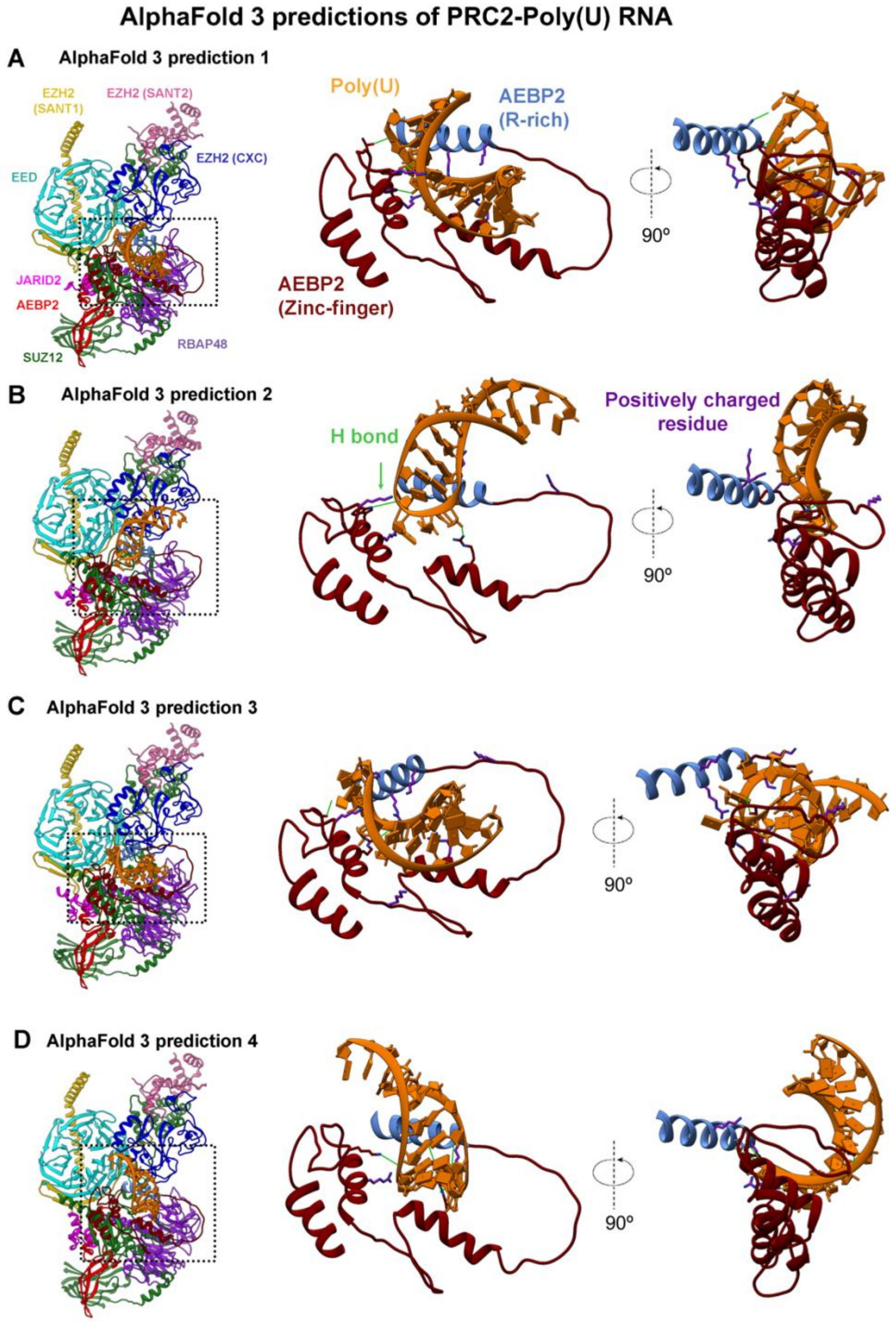


**Figure S9. AlphaFold 3 predicted interactions between AEBP2 and Poly(U) RNA.** (A-D) Left: Overall models from AlphaFold 3 predictions. Dashed boxes are zoomed in. Center and Right: Zoom-in views of $TERRA_{mut}$ RNA, AEBP2 arginine-rich segment, and zinc-finger domains. Hydrogen bonds were analyzed by the FindHbond function built in Chimera and highlighted in green color. Side chains of positively charged residues aiming towards RNA are also highlighted in purple color.

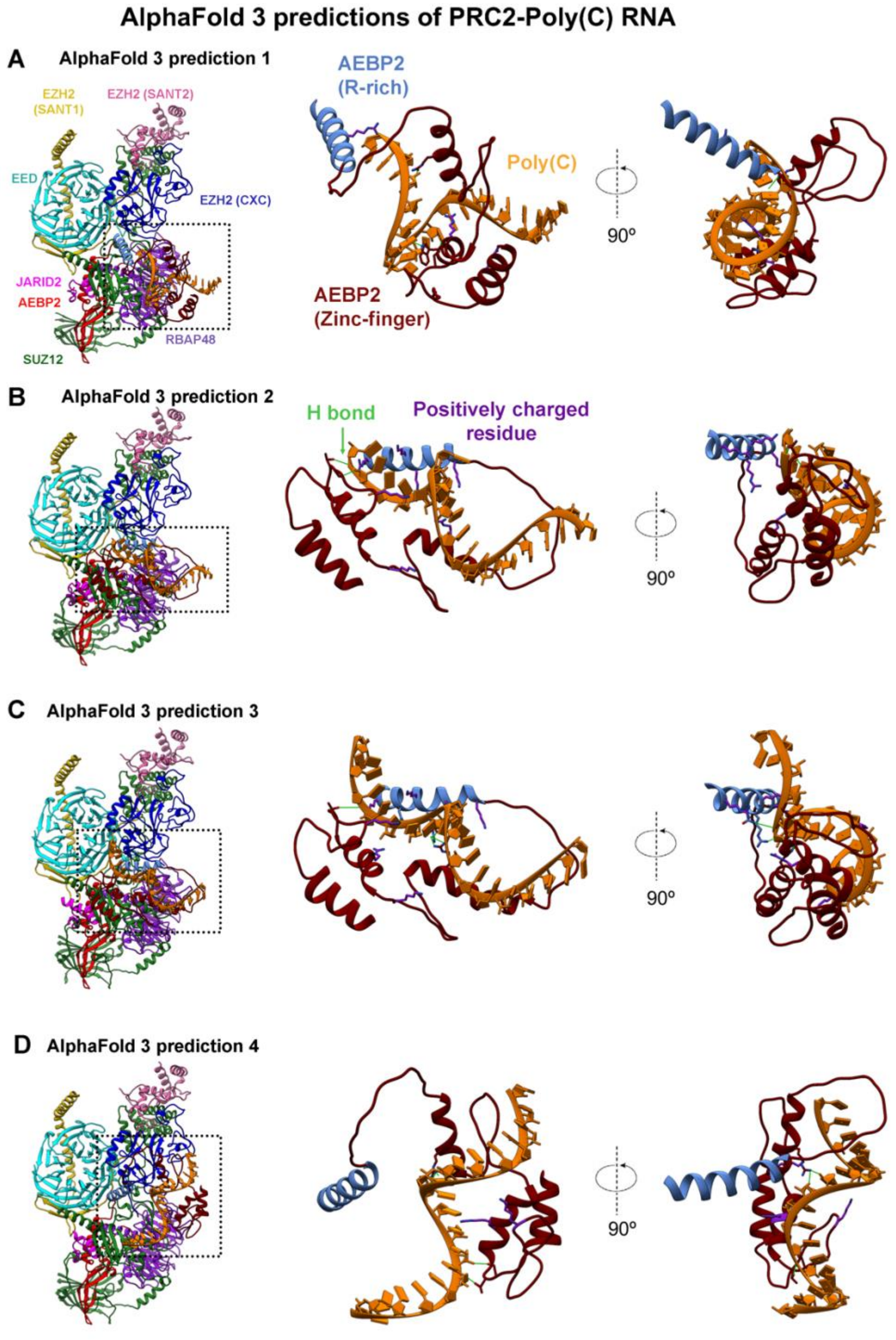


**Figure S10. AlphaFold 3 predicted interactions between AEBP2 and Poly(C) RNA.** (A-D) Left: Overall models from AlphaFold 3 predictions. Dashed boxes are zoomed in. Center and Right: Zoom-in views of $TERRA_{mut}$ RNA, AEBP2 arginine-rich segment, and zinc-finger domains. Hydrogen bonds were analyzed by the FindHbond function built in Chimera and highlighted in green color. Side chains of positively charged residues aiming towards RNA are also highlighted in purple color.

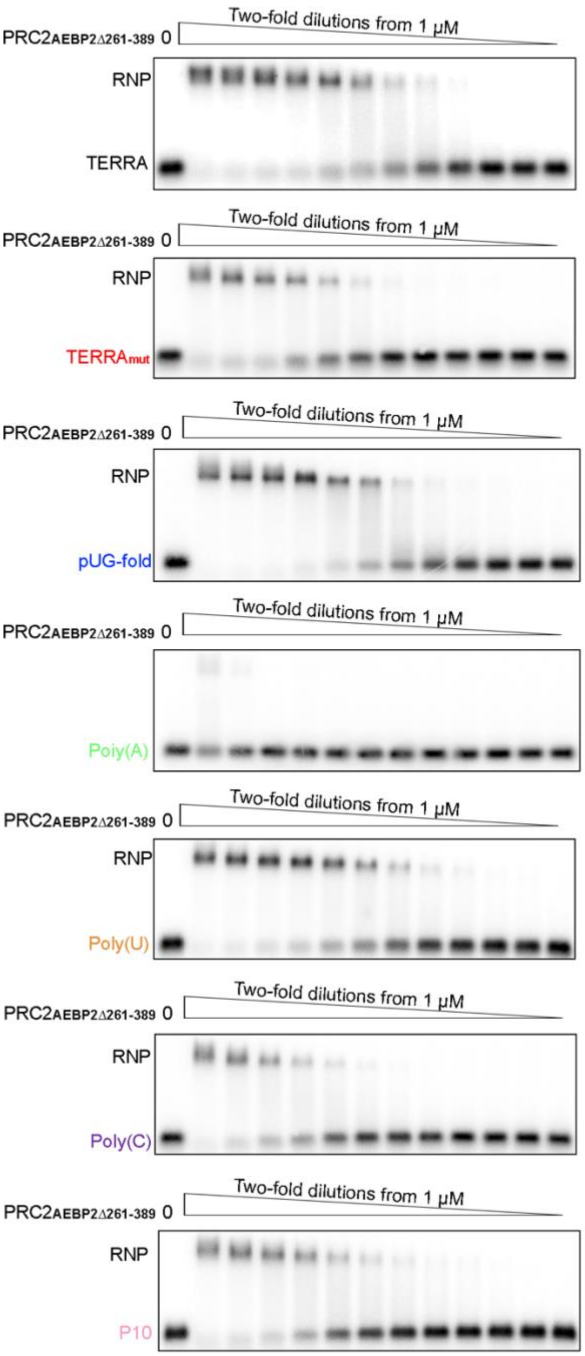


**Figure S11. PRC2$_{AEBP2\Delta 261\text{-}389}$ mutant has decreased binding towards TERRA$_{mut}$ and Poly(C).** Representative EMSA gels. Quantification of EMSA results are shown in Figure 3C.

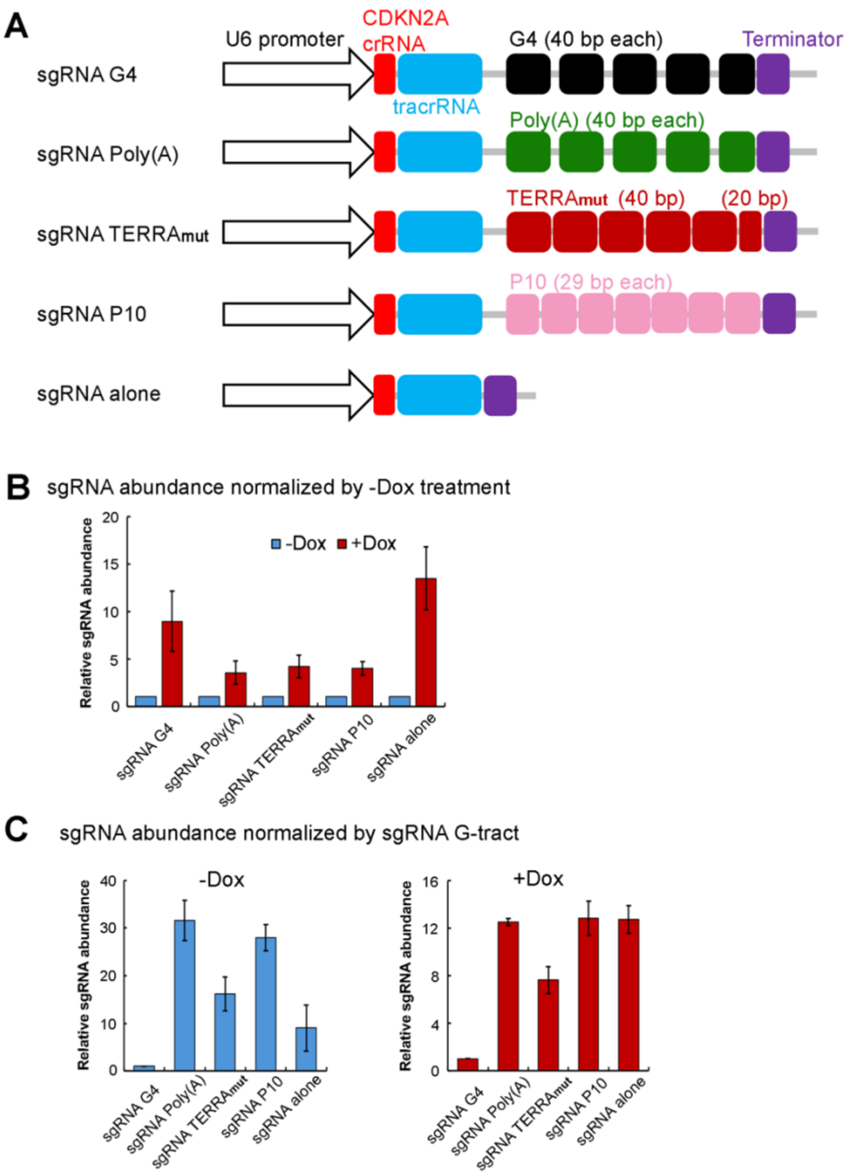


**Figure S12. Adapted CRISPR Display strategy to examine the ability of different RNA elements to regulate PRC2 in cells.** (A) Schematic of five modified sgRNA constructs used in this study. (B) qRT-PCR analyses measure the relative abundance of each sgRNA normalized by the -Dox treatment of the same RNA. This result indicates that the expression of dCas9 protein by Dox induction can stabilize sgRNA independent of the RNA appended to the tracrRNA, suggesting the formation of dCas9-sgRNA RNP in all five cell lines. (C) qRT-PCR analyses measure the relative abundance of sgRNA normalized by sgRNA G-tract in -Dox treatment (left) and +Dox treatment (right) separately. sgRNA G-tract is significantly less abundant compared to other sgRNAs, which could be due to reduced expression or lower RNA stability in cells.

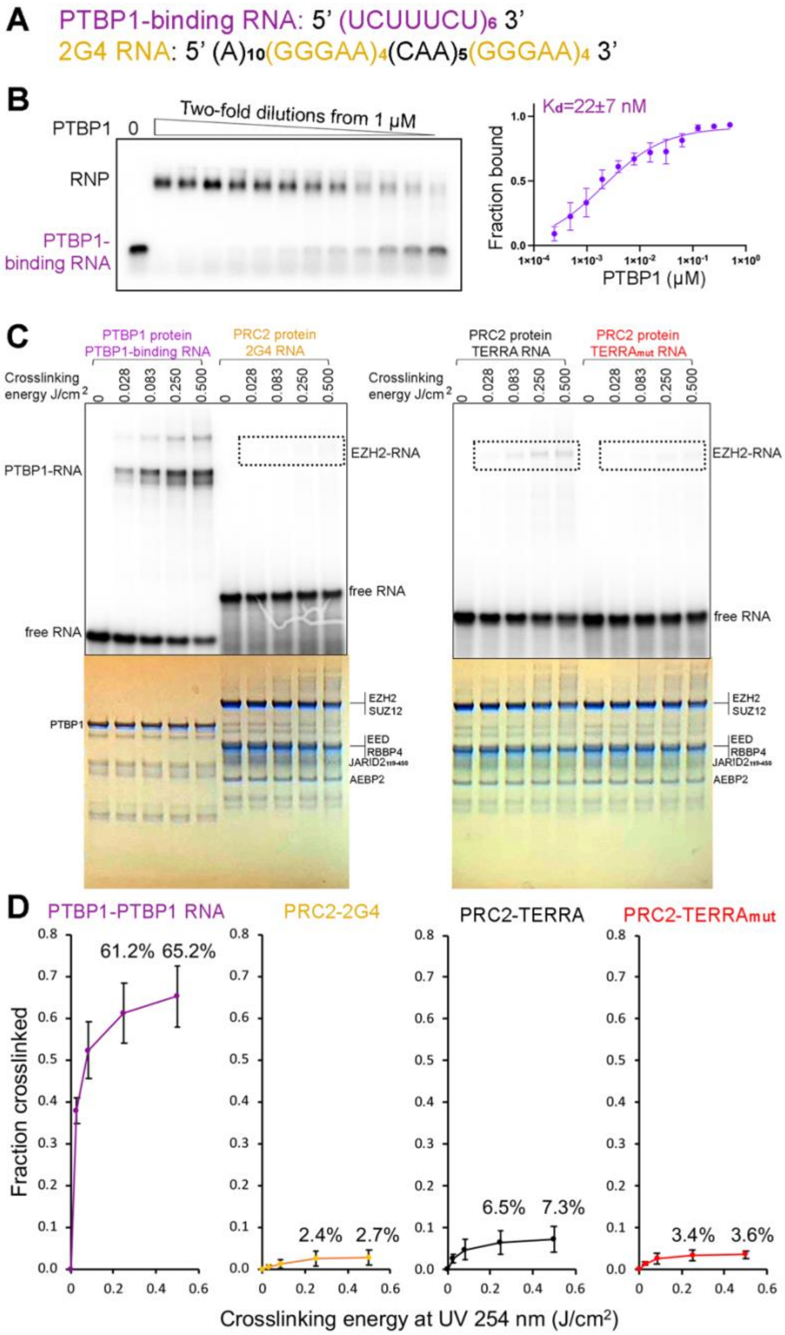


**Figure S13. UV crosslinking has very low efficiency for capturing PRC2-RNA associations.** (A) Sequences of RNA oligos used in this assay. (B) EMSA result of recombinant PTBP1 binding to $(UCUUUCU)_6$ RNA. Left: representative EMSA gel. Right: quantification of EMSA results. Error bars are range of two replicates. (C) Representative UV crosslinking assays. In all reactions, proteins were 0.6 μM (at least 10-times $K_d$) to ensure saturated RNA-binding. Gels were imaged to reveal crosslinking of radiolabeled RNA (top) or stained with Coomassie blue to confirm equal loading of proteins (bottom). Dashed boxes are positions expected for crosslinked EZH2-RNA complexes. (D) Quantification of three UV crosslinking replicates. Error bars are mean ± standard deviation.

**Table S1. Cryo-EM data collection and refinement statistics**

| | Multibody body 1 | Multibody body 2 | Consensus map |
|---|---|---|---|
| **Data collection and processing** | | | |
| Magnification | 130,000 | 130,000 | 130,000 |
| Voltage (kV) | 300 | 300 | 300 |
| Camera | Falcon 4 | Falcon 4 | Falcon 4 |
| Electron exposure ($e^-/Å^2$) | 50 | 50 | 50 |
| Exposure rate (e-/pixel*s) | 8.78 | 8.78 | 8.78 |
| Number of frames | 1323 | 1323 | 1323 |
| Defocus range (μm) | -0.5 to -1.9 | -0.5 to -1.9 | -0.5 to -1.9 |
| Pixel size (Å) | 0.97 | 0.97 | 0.97 |
| Symmetry imposed | C1 | C1 | C1 |
| Movies collected (no.) | 13,028 + 1202 (tilted stage) | 13,028 + 1202 (tilted stage) | 13,028 + 1202 (tilted stage) |
| Initial particle images (no.) | 3,194,364 + 191,127 (tilted stage) | 3,194,364 + 191,127 (tilted stage) | 3,194,364 + 191,127 (tilted stage) |
| Final particle images (no.) | 105,974 + 14,684 (tilted stage) | 105,974 + 14,684 (tilted stage) | 105,974 + 14,684 (tilted stage) |
| **Map resolution (Å)** | | | |
| FSC0.143 (unmasked/masked) | 3.8/3.3 | 3.8/3.4 | 3.7/3.4 |
| Map resolution range (Å) | 3-10 | 3-11 | 3-11 |
| **Refinement** | | | |
| Initial model used (PDB code) | 8FYH | 8FYH | 8FYH |
| Resolution cutoff (Å) | 4 | 4 | 4 |
| Map sharpening *B* factor ($Å^2$) | 0 | 0 | 0 |
| **Model composition** | | | |
| Non-hydrogen atoms | 12111 | 12103 | 24433 |
| Protein residues | 1793 | 1793 | 3586 |
| Nucleotide residues | 0 | 0 | 10 |
| Ligands | ZN:7 | ZN:7 | ZN:14 |
| ***B* factors ($Å^2$)** | | | |
| Protein | 24.35 | 13.42 | 13.42 |
| Ligand | 44.27 | 22.70 | 22.70 |
| **RMSD** | | | |
| Bond lengths (Å) | 0.002 (0) | 0.002 (0) | 0.004 (0) |
| Bond angles (°) | 0.521 (4) | 0.536 (6) | 0.971 (4) |
| **Validation** | | | |
| MolProbity score | 2.25 | 2.27 | 2.26 |
| Clashscore | 9.77 | 8.81 | 9.57 |
| Poor rotamers (%) | 2.01 | 2.58 | 2.18 |
| **Ramachandran plot** | | | |
| Favored (%) | 91.43 | 92.12 | 91.66 |
| Allowed (%) | 8.45 | 7.77 | 8.17 |
| Disallowed (%) | 0.11 | 0.11 | 0.17 |

**SUPPLEMENTARY DOCUMENT REFERENCE**

### Bibliography and References Cited

1. Monchaud, D. (2023). Why does the pUG tail curl? Mol Cell *83*, 330-331. 10.1016/j.molcel.2022.12.010.
